# COLLEMBOT: AI-Based Counting of Collembola for OECD 232 Tests

**DOI:** 10.64898/2026.01.16.697653

**Authors:** Micha Wehrli, Adrian Meyer, Éverton Souza da Silva, Sam van Loon, Bart G. van Hall, Cornelis A.M. van Gestel, Tiago Natal-da-Luz, Max V.R. Döring, Heike Feldhaar, Magdalena Mair, Denis Jordan, Miriam Langer

## Abstract

Ecotoxicological tests with soil organisms, such as the collembolan *Folsomia candida*, are essential for assessing chemical risks in terrestrial ecosystems. However, the current Organization for Economic Co-operation and Development (OECD) 232 reproduction tests rely on manual counting of juvenile and adult Collembola, a process that is costly, labor-intensive, time-consuming and prone to operator bias. These limitations restrict data availability and hinder robust risk assessments.

We therefore developed COLLEMBOT, an automated counting tool based on a YOLOv11 convolutional neural network, designed to integrate seamlessly into OECD workflows without protocol modifications. The model was trained on high-resolution images (n = 3207) from multiple laboratories and validated using 22 independent datasets (n = 1704 images) from Amsterdam (Netherlands), Basel (Switzerland), Bayreuth (Germany), Coimbra (Portugal) and Aarhus (Denmark). Datasets consisted of relevant standard soils (OECD artificial soils with 2.5%, 5% and 10% sphagnum peat; LUFA 2.2) and the springtail *Folsomia candida.* Automated counts showed strong agreement with manual counts (R² = 0.88–0.99). Dose-response curves derived from automated and manual counts strongly overlapped and effect concentrations (EC₁₀ and EC₅₀) differed minimally (Median %Δ 6.2 ± 23 and EC_10_ - EC_90_ R^2^ ≥ 0.977), remaining within acceptable limits for regulatory risk assessment and confirming reliability.

Time efficiency improved significantly: a test with ∼300 images and up to 1,500 individuals per image was processed in less than 3 hours, compared to ∼137 hours needed for manual counting, a reduction of approximately 97%. By reducing labor and improving reproducibility, COLLEMBOT enables broader hazard data generation for collembolans, supporting science-based chemical risk assessment. The code and workflow are publicly available to facilitate adoption and community-driven development.

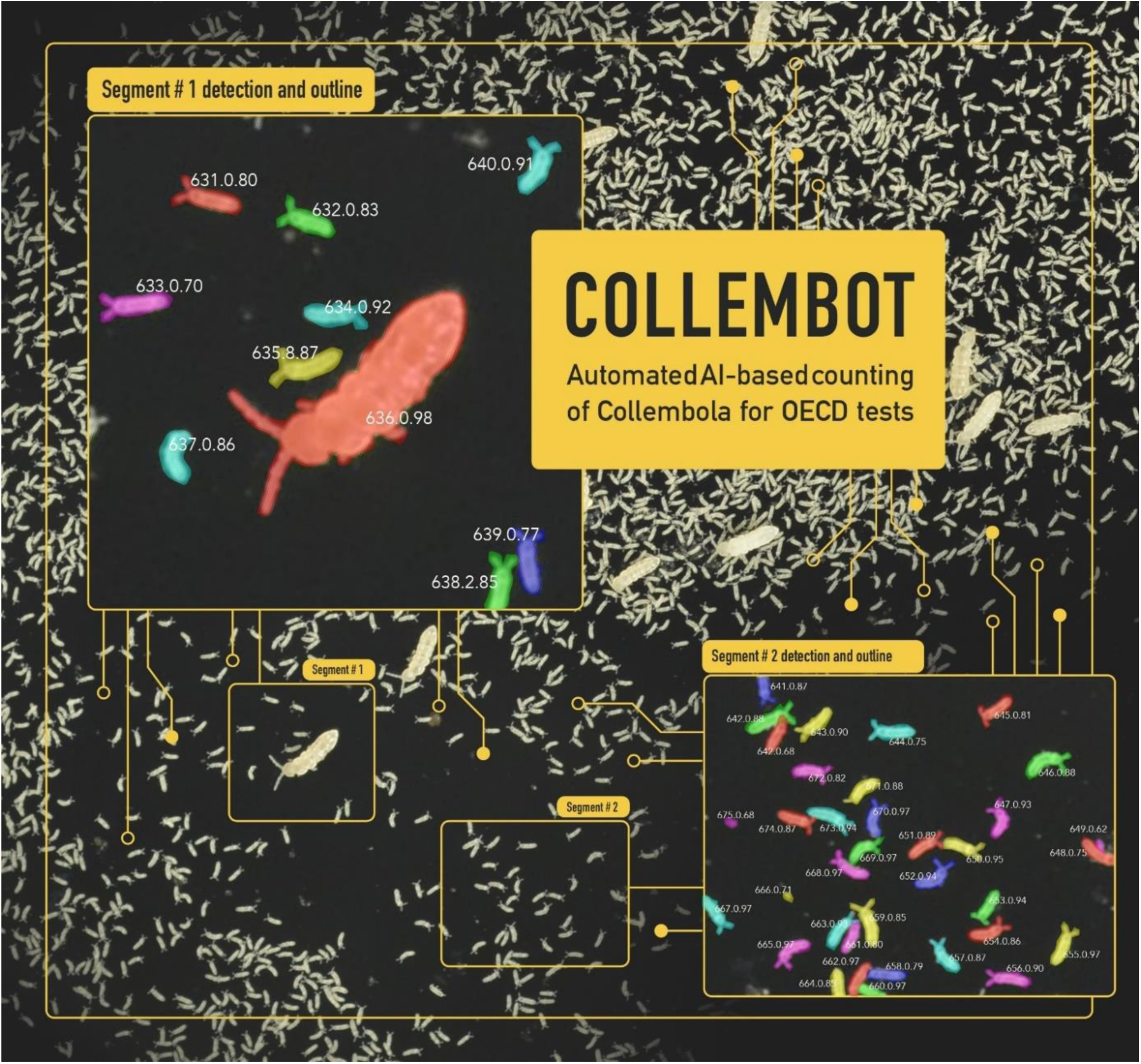

## Introduction

The number of chemicals produced and applied has increased drastically (Z. Wang et al., 2020). Some of these chemicals will, one way or another, enter the environment and pose a risk to biodiversity. Thus, it is essential to conduct (holistic) pro- and retrospective approaches to estimate the risk of chemicals (e.g., novel substances and substances of emerging concern) and visualize existing risks (i.e. substances already on the market). Currently, the hazard data requirements differ depending on the jurisdiction and the chemical’s intended use case: industrial chemicals (in Europe: Registration, Evaluation, Authorization and Restriction of Chemicals (REACH)), biocides, pharmaceuticals, or plant protection products, each with different regulations and hazard data requests. To date only a few chemical regulations require toxicity data for soil organisms, for instance the registration of chemicals for agricultural use (e.g., Regulation 1107/2009 in Europe (EC, 2009)). Despite the high cost of acquiring these data, they are necessary to assess the risk of an increasing number and / or concentrations of substances. In addition to the huge number of chemicals, particulate pollutants, including microplastics, have recently gotten increasing attention due to their negative effects on various organisms (Hampton et al., 2025; Lead et al., 2018; van Loon et al., 2025). As particulate pollutants tend to accumulate particularly in soil, they are expected to become notably problematic for soil dwelling organisms (De Souza Machado et al., 2018; Lead et al., 2018). To ensure that thorough and broad risk assessment, for both already known pollutants and emerging contaminants of concern, is not hindered by financial costs for hazard assessments, we need a streamlined and robust testing approach to generate hazard data.

The risk assessment for chemicals in soil is based on standardized laboratory toxicity tests, such as the Organization for Economic Co-Operation and Development (OECD) test guidelines 232/226/222 (OECD, 2016a, 2016b, 2016c), with a primary focus on reproductive effects on the tested organisms. The primary animals for these tests are earthworms (*Eisenia fetida* or *Eisenia andrei*), predatory mites (*Hypoaspis aculeifer*), and springtails (*Folsomia candida*), respectively. These tests are usually conducted over 56, 14 and 28 days for earthworms, mites and springtails, respectively, and do not require much labor during the exposure phase. However, when these tests are complete, each test jar contains a substantial number of juveniles, which need to be counted. In particular for the springtail *Folsomia candida,* the number of juveniles can exceed 1000 individuals in one test jar (Krogh et al., 2008) which, when manually counted, can add up to many hours of repetitive work and is prone to operator errors.

After extracting the springtails by heat extraction or flotation, the counting can be performed either by eye or with a microscope directly in the extract, or by photographing them to later count them by using, for instance, digital image tools. The most commonly used tool for these assays is ImageJ (Schneider et al., 2012). ImageJ and similar software allow users to mark individuals manually or apply threshold-based automated counting to highlight the animals. However, these approaches are still labor-intensive and time-consuming either placing markers for each individual or adjusting contrast and threshold settings for every image. This makes the process slow, leading to a high number of repetitive working hours and is highly subjective to the operator and can thus lead to observer bias in the counting process (Abreu et al., 2022), especially when hundreds of images with varying lighting and background conditions must be processed. Threshold-based automated counting and batch processing is also limited because springtails often overlap or cluster, requiring manual correction to avoid miscounts.

Similar problems have been observed in other standardized tests including for instance tests on the water flea *Daphnia magna* (Abreu et al., 2022) and the potworm *Enchytraeus crypticus* (van Hall & van Gestel, 2025), or for the OECD micronucleus tests (Xue et al., 2025). In several cases, the development of an automated counting source sped up the counting process, as demonstrated by Xue et al. (2025) who reported a 20x increase in hourly counting speed and up to 60x in daily counting speed. Van Hall & Van Gestel (2025) showed that manual counting of 1620 pictures with enchytraeids took approximately 270 hours, while automated counting reduced this time to approximately 6.75 hours. In the case of springtail counting, the working time saved could be substantial. A manual count of one test picture can take up to 60 minutes, depending on the number of springtails per image, the quality of the image and the person who is counting them. Hence, there is a need and, obviously, potential in automating the counting for OECD springtail toxicity tests.

Previous studies applied different approaches to automated image analysis, however, these had drawbacks when applied to OECD 232 standard testing and require minor to major modifications to extraction protocols, which complicates the process and increases the use of materials and time. For instance, the use of anesthesia tools (e.g., chill coma or ethanol) and thermal imaging (Pang et al., 2023), or relying on a special device and low-density moving collembolans is not suitable for high-throughput OECD tests (Bánszegi et al., 2014). The method described in the OECD 232 guideline using contrast-enhanced counting in ImageJ (Caridade et al., 2011; Krogh et al., 1998) relies heavily on manual labor and is not as reliable and versatile as computer vision, which, in turn, has become more robust in recent years. Computer vision refers to advanced algorithms, often based on deep learning, that enable machines to interpret and analyze visual information in a way that mimics human perception, whereas ImageJ relies on rule-based thresholding and segmentation for object detection, which is less adaptable and robust to variations in lighting, background, and object morphology. In 2022, Sys and colleagues (Sys et al., 2022) developed a new model that utilizes computer vision to identify different Collembola species in fluid samples and already achieved a good F1 score of 94%. The F1 score is a standard metric that reflects the overall accuracy of detection by combining both precision and recall. Although the approach seems promising, the study was focused on counting collembolans preserved in fluid and not on soil surfaces. A similar approach was proposed by Oriol and colleagues who optimized a model to seek out Collembola on a larger scale, but it was not optimized for OECD testing (Oriol et al., 2024).

These examples highlight that, despite the existence of several approaches for automating counting in ecotoxicological tests there is currently no standardized, widely applicable and GLP-compliant solution for OECD 232 springtail tests. Existing methods either require protocol modifications, additional materials, or specialized equipment, which limits their scalability and regulatory acceptance. Furthermore, most approaches fail to achieve sufficient accuracy and robustness under varied imaging conditions, making them unsuitable for routine risk assessment workflows. This underscores the need for a method that combines high accuracy, adaptability to different laboratory setups, and compatibility with existing OECD guidelines without introducing additional complexity or cost.

The goal of this study was to develop an automated springtail counting tool, COLLEMBOT, that can speed up the quantification of the absolute number of *Folsomia candida* (or other springtail species) in standard OECD 232 toxicity tests, with a reliable and robust output that is comparable or as good as a human but lacking the operator bias, higher comparability and with a significant decrease of processing time. It also aimed at making COLLEMBOT eligible for validation to comply with GLP requirements. To achieve this and facilitate the applicability to different laboratory setups, image data from 20 independent previously counted OECD tests with different soil types and taken with different camera devices in 5 different labs were used for model training. The performance of the trained model was assessed using several independent test datasets that were excluded from the training process and model optimization. The developed model is accurate and consistent under varied imaging conditions and is made publicly accessible for use and future adjustments.

## Material & Methods

For the published test sets, Aarhus University, Denmark (Wehrli et al., 2024) and Vrije Universiteit Amsterdam, Netherlands (van Hall et al., 2025) as well as the training set CollembolAI (Sys et al., 2022), the organisms and test set ups are described in the relative source. The organisms and test setups of unpublished test sets can be found in Supplementary Information S1. An overview of the different test sets is given in Supplementary Table 1.

### Test pictures

Test pictures to train the model were taken from 5 different laboratories (FHNW Basel, Vrije Universiteit Amsterdam, University of Bayreuth, Cloverstrategy Lda Coimbra, and a dataset from CollembolaAI (Sys et al., 2022)). The non-OECD image dataset from CollembolaAI was added to increase the number of annotated tiles and increase the variation and, thus, robustness of the model through image pyramids, further description in the section multiscale datasets below.

Different test soils were used, including the natural LUFA 2.2 soil and different OECD artificial standard soils with different organic matter contents (OM; added as sphagnum peat) (OECD 2.5% OM, 5% OM, 10% OM). Additionally, the Bayreuth training dataset contained microplastic particles in the soil (polystyrene, 0.5% w/w, size range of the fragments between 20-75 µm). These pictures originated from different OECD 232 tests with *Folsomia candida,* kept in different test soils and taken with different camera set ups. Furthermore, at the end of the experiments, after adding water to the test jars for the springtails to float, a foam cover of varying density developed on the surface (see Figure 1). Such foam is naturally formed, and, in the context of these image datasets, it can increase the complexity of the surface and, consequently, the model. These datasets were then fed through the pipeline to generate labels on the pictures, which were later manually corrected in labelme (Russell et al., 2008) to obtain a correct ground truth. An overview of all datasets, including laboratory origin, soil type, chemicals tested, camera setup, image quality and purpose (training vs validation), is provided in Supplementary Table 1 and described in detail in Supplementary Information S1 An overview of the heterogenicity of the samples can be seen in figure 1.

**Figure 1.**
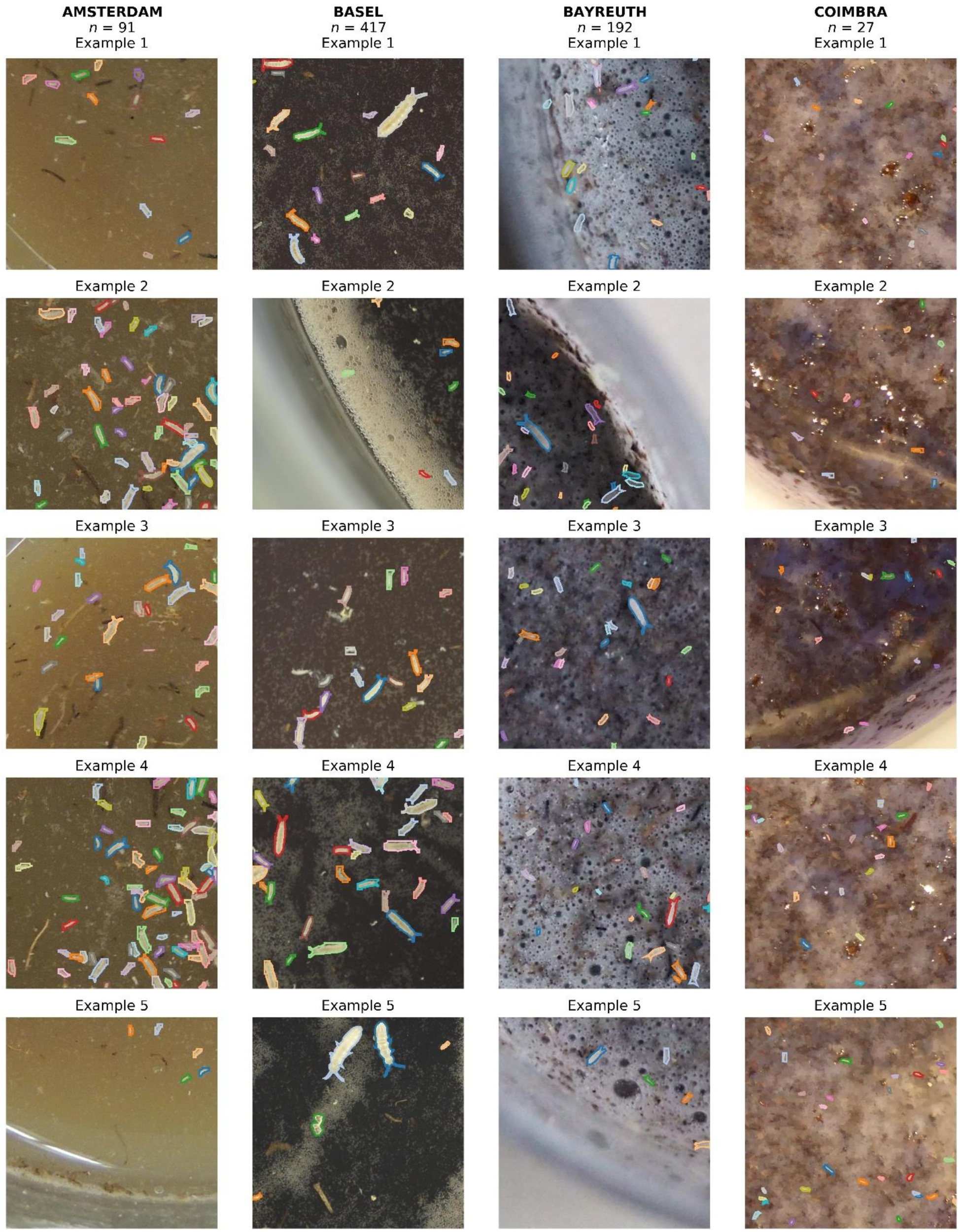
Representative image tiles from high-resolution datasets collected during Collembola (springtail) toxicity tests. The images illustrate differences in image quality, contrast, and substrate conditions across four laboratories (Amsterdam, Basel, Bayreuth, and Coimbra). Each tile shows examples of test jars with Collembola individuals (colored overlays indicate the detected organisms) used for automated counting in eco-toxicological assays. Variations in lighting, background texture, and substrate composition highlight challenges for image-based quantification of springtail survival and reproduction.

Additional 24 sets of pictures (Vrije Universiteit Amsterdam (van Hall et al., 2025), FHNW Basel (Unpublished), University of Bayreuth (Unpublished), Cloverstrategy Lda Coimbra (Unpublished) and Aarhus University (Wehrli et al., 2024)) with already hand counted springtails were used to validate the model’s capability to count similar numbers as a human operator would.

### Model Training

### Multiscale Datasets

In addition to the training datasets from OECD collembolan tests, we tested whether extending our datasets with multiscale image pyramids of the Collembola AI dataset (Sys et al., 2022) (Figure 2) would increase robustness of model training and evaluation across various object scales and densities. The original images (Level 1(L1)) exhibited high-resolution scans with clearly defined Collembola individuals on a relatively uniform black background. Here, L1–L16 denote the levels of the multiscale image pyramid, where L1 corresponds to the original high-resolution tiles and higher levels (e.g., L2, L4, L8, L16) represent progressively coarser resolutions created by merging tiles from the previous level, increasing the field of view and organism density. The direct usage of highest resolution image tiles resulted in an overrepresentation of very large Collembola segments. However, to generalize model performance, coarser-resolution levels (L2 n=1745, L4 n=555, L8 n=144, L16 n=36; n = number of images in Level) were introduced by progressively combining multiple tiles from the previous finer resolution.

**Figure 2.**
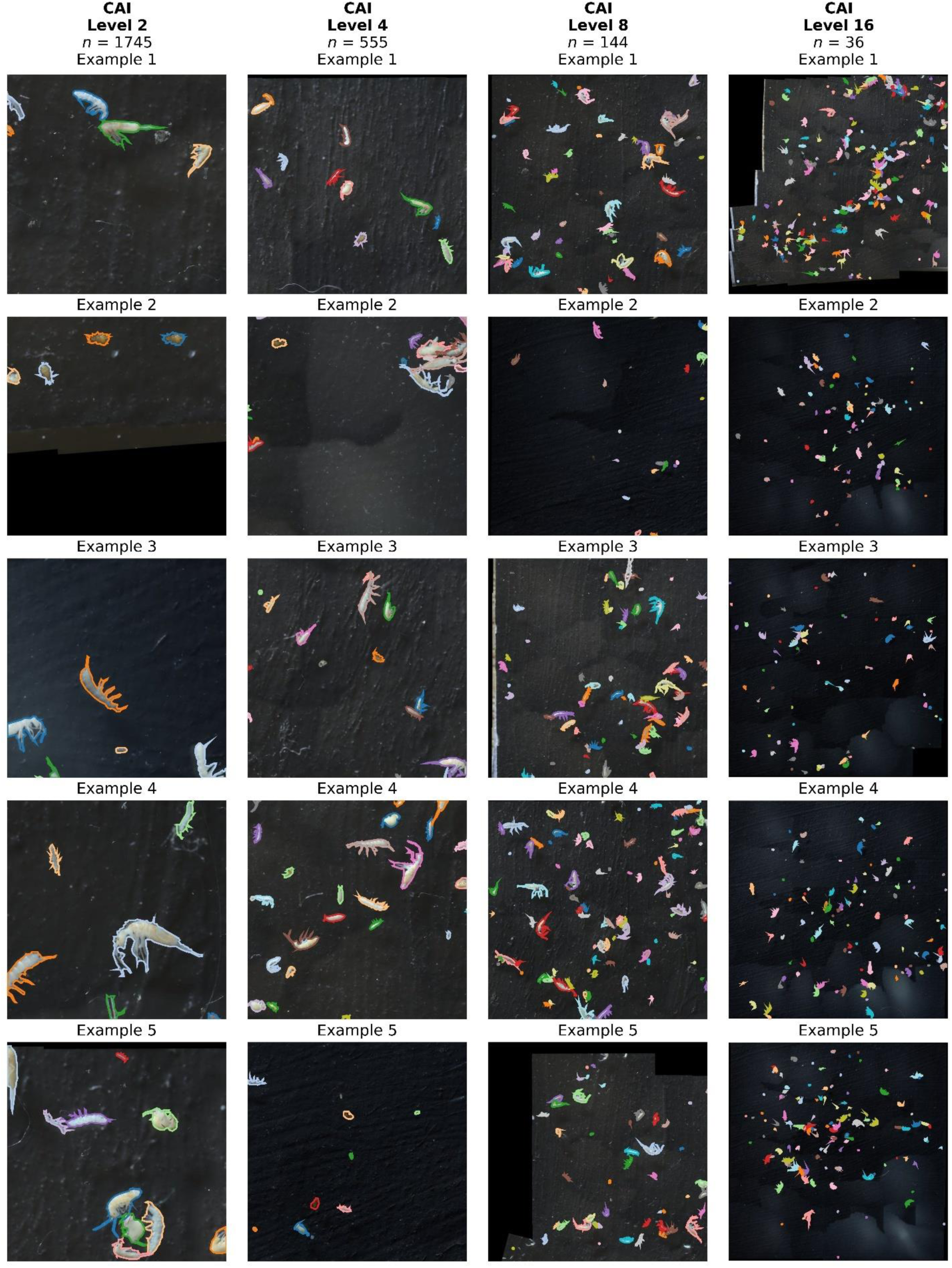
Examples of multiscale image pyramid levels (L2–L16) adapted from the Collembola AI dataset (Sys et al., 2022), containing 12 different springtail species. Each column represents a different pyramid level, where higher levels correspond to progressively coarser resolutions and larger fields of view, resulting in increased organism density per image. This multiscale approach was used to improve model robustness across varying object sizes and densities.

This multiscale strategy aims to simulate practical scenarios where object size and image resolution vary considerably, thus enhancing the models’ ability to maintain high detection and segmentation accuracy regardless of scale. The pyramid structure also systematically represents increasingly dense distributions of Collembola, improving the robustness of predictions under varying organism density and ensuring more reliable applicability to real-world datasets.

### Model Training

Model training was carried out on an HP Apollo high–performance computing (HPC) system equipped with 4 × NVIDIA Tesla V100 GPUs (32 GB VRAM each) under Linux. Training and inference used a Python 3.10 stack with Conda for environment management, PyTorch with CUDA support for Deep Learning, the Ultralytics implementation for YOLO v11–xl–seg (Khanam & Hussain, 2024; Redmon et al., 2016) and a detectron2 implementation for Mask R–CNN (He et al., 2018).

YOLO v11–xl segmentation (YOLO v11–xl–seg) was used as the primary model in the pipeline and for all automated counts reported here. Mask R–CNN was trained on the same data as a comparative baseline. Both models were configured to detect a single biological class (“Collembola”). During preprocessing, all annotation labels referring to collembolans were mapped to this class.

Images were annotated in LabelMe with polygon outlines around individual Collembola. A custom preprocessing script collects all image-annotation pairs from the different source datasets, sanitizes their naming and converts the LabelMe polygons into the input formats required by YOLO v11–xl–seg and Mask R–CNN (normalized polygon coordinates with a single class label). To obtain reproducible data splits, each source dataset was randomly divided into training, validation and test images using fixed proportions (70/15/15 %). For every dataset, the test images were written to a small “reserve file”; comparative evaluations reuse this exact test set, ensuring that test results are directly comparable between models and experiments.

YOLO11x–seg (Release 8.3.0) was initialized from publicly available standard COCO-pre–trained weights (transfer learning) and fine–tuned on our data. The main hyperparameters (image size, batch size, number of epochs, learning rate schedule, data augmentation and GPU devices) were defined in a YAML configuration file and kept constant across experiments and optimized leveraging automated Optuna studies (Akiba et al., 2019). During training, the framework recorded standard detection metrics (precision, recall, F1 score, mean average precision) per epoch and saved both the best and final model weights.

Mask R–CNN was trained on the same training and validation images and annotations, also using a COCO–pre–trained backbone and comparable optimization settings (number of epochs, batch size, input resolution). Mask R–CNN results are used only for comparison; the production pipeline described below uses YOLO v11–xl–seg. Both models were evaluated on the held–out and independent test sets with additionally computed dataset–specific metrics (one test evaluation per source dataset) to check robustness across imaging conditions.

### Inference

For local inference use of the trained YOLO model, a single CUDA–capable GPU with ≥ 8 GB VRAM (16 GB recommended) and ≥ 16 GB system RAM on a Unix–like operating system (Linux or Windows via WSL2) is sufficient. The pipeline can alternatively be executed on a cloud GPU environment (e.g. Google Colab).

### Post–processing: tiling and fusion of detections

To restrict the analysis to the circular region of interest, we first automatically detected the main round surface in each image using the Hough circle transform (Ioannou et al., 1999). For every slide, the original Red, Green, Blue (RGB) image was loaded and converted to grayscale, then smoothed with a 5×5 median filter to reduce noise while preserving edges. We applied OpenCV’s gradient-based *Hough-Circles* function to the preprocessed image, with the accumulator resolution set to 1.2 relative to the image resolution and the minimum distance between candidate circles set to one quarter of the image height. To ensure that we detected only the large circular sample area (e.g. the dish or slide region), we restricted the search radius to be between 20% and 49% of the image height. When multiple circles were found, we selected the one whose center was closest to the image center, under the assumption that the sample surface was approximately centered in the field of view. This detected circle was then used as a mask: for each predicted object polygon, we computed its centroid and retained only those polygons whose centroids laid inside the circle, discarding all detections outside the region of interest. If no valid circle was detected, the masking step was skipped, and all polygons were kept. This procedure effectively limits subsequent fusion and counting to the relevant circular surface, reducing spurious detections from the image background.

The original images were large and could not be reliably processed at once at high resolution. We therefore applied YOLO v11–xl–seg on smaller image tiles. Each image was split into square tiles of 576 × 576 pixels. To reduce edge effects, we used two overlapping tiling grids: one without offset and one shifted by half a tile in both directions. All tiles were processed by the trained YOLO v11–xl–seg model on multiple GPUs in parallel inference. For each tile, the model provided instance masks and confidence scores for all detected collembolans. Tile–level detections were then mapped back to global image coordinates by adding the tile offsets, yielding a set of partially overlapping polygons per original image. This produced a “raw” set of candidate detections for each image, often with duplicates near tile borders. These raw polygons were subsequently merged into final objects by a fusion step.

In the final pipeline we used a graph–based fusion (“graphcut”) approach (Boykov & Jolly, 2001): Overlapping polygons with an intersection–over–union (IoU) above a threshold were grouped into clusters. For each cluster, all polygon boundaries were combined into a single smooth object outline using a concave–hull (alpha–shape) construction. The confidence score of the fused object was taken as the highest confidence among its constituent polygons under the assumption that a completely visible hexapod will yield higher confidence scores than a partially visible section. This fusion reduced multiple overlapping tile–level detections to one biologically meaningful object per collembolan individual. As an additional advantage of this method, the convex outer shape approximation of each individual was retained.

Several alternative strategies for merging overlapping detections were explored in addition to the graph–based fusion described above, including:

- Simple non–maximum suppression (keeping only the highest–confidence polygon among overlapping candidates),
- Union–based fusion (geometric union of overlapping polygons),
- “Voting” approaches that rasterize and average multiple overlapping masks before contour extraction, and
- Tile–boundary heuristics that explicitly favor detections not clipped by tile edges.

To select the most suitable strategy and its parameters, we conducted a hyperparameter optimization using Optuna on a subset of images with manual annotations. In this optimization, the fusion method, the detection of a confidence threshold and the IoU threshold were varied, and performance was quantified by the mean F1–score (harmonic mean of precision and recall) across these images. The study consistently identified the ”graphcut” approach as the best compromise between missing individuals and over–segmentation, and the optimized settings were adopted for all subsequent analyses.

### Comparison of hand–counted and automated counts

#### Model Validation of training validation datasets

To assess how well the automated method reproduced manual counts, we compared YOLO v11–xl–seg outputs with hand–counted collembolan numbers. For each image of the training datasets with manual annotations, the centroid coordinates of all manually marked individuals were available as CSV files. Automated detections from the fused YOLO output were treated as polygons and manual points were classified as: (i) True positives (TP) if they fell inside any polygon, (ii) False negatives (FN) if they were not covered by any polygon, (iii) polygons not containing any manual point were counted as false positives. For every image we calculated precision, recall and F1–score, as well as the total number of individuals counted manually and automatically. Across all manually annotated images, we then computed overall precision, recall and F1–score, and calculated the coefficient of determination (R²) between manual and automated counts using linear regression. This provided a detection-response–type assessment of how reliably the automated pipeline reproduced manual collembolan counts under the conditions of the ecotoxicity tests.

#### Statistical Analysis on independent datasets

Model validation under real test conditions was performed using previously published datasets (See Table Supplementary, Table 1) from Amsterdam (n= 1281 pictures) (van Hall et al., 2025), Bayreuth (n= 56 pictures) (unpublished), Basel (n=123 pictures) (unpublished) and Denmark (n= 209 pictures) (Wehrli et al., 2024), Coimbra (n= 35 pictures) (unpublished), which were not used for training. These datasets comprised four different test soils (LUFA 2.2, OECD 2.5, 5 and 10). Automated counts generated by the YOLO-based pipeline were compared to manual counts through three approaches, linear regression and dose-response modelling is described here, while the NOEC approach is described in Supplementary Information S5.

#### Linear Regression

Agreement between manual and automated counts was assessed using simple linear models (lm(Automated ∼ Manual)), reporting slope, intercept and coefficient of determination (R²) for each substance.

#### Dose–Response Modeling

Both manual and automated datasets were fitted with the same nonlinear model type (three-parameter Weibull, W1.3), following OECD Guideline 232. Analyses were conducted in R (version R version 4.4.2) using the drc package (Ritz et al., 2015) within RStudio (Posit team, 2025). Effect concentrations (EC_10_, EC_20_, EC_50_, EC_90_) and 95% confidence intervals were calculated via the delta method using the ED() function.

All concentration-response data and statistical outputs were summarized in Supplementary Information S4 tables 3-5, including NOEC, LOEC and the type of test applied.

Observed means ± standard errors per concentration were summarized and visualized alongside predicted curves using ggplot2 (Wickham, 2016). Comparative ED estimates were tabulated and plotted as point–error bar charts across substances and effect levels. All scripts ensured reproducibility by applying consistent model structures and confidence interval estimation for both manual and automated datasets.

## Results

### Model Selection

The comparative evaluation of Mask R-CNN and YOLO v11 segXL across the five imaging setups (Figure 3) reveals systematic differences in detection performance that depended both on image quality and on the evaluation metric considered. To provide a balanced assessment, we jointly report mean average precision at IoU 0.5 (mAP50) and F1-scores for bounding box (BBox) detection and instance segmentation (Mask).

**Figure 3.**
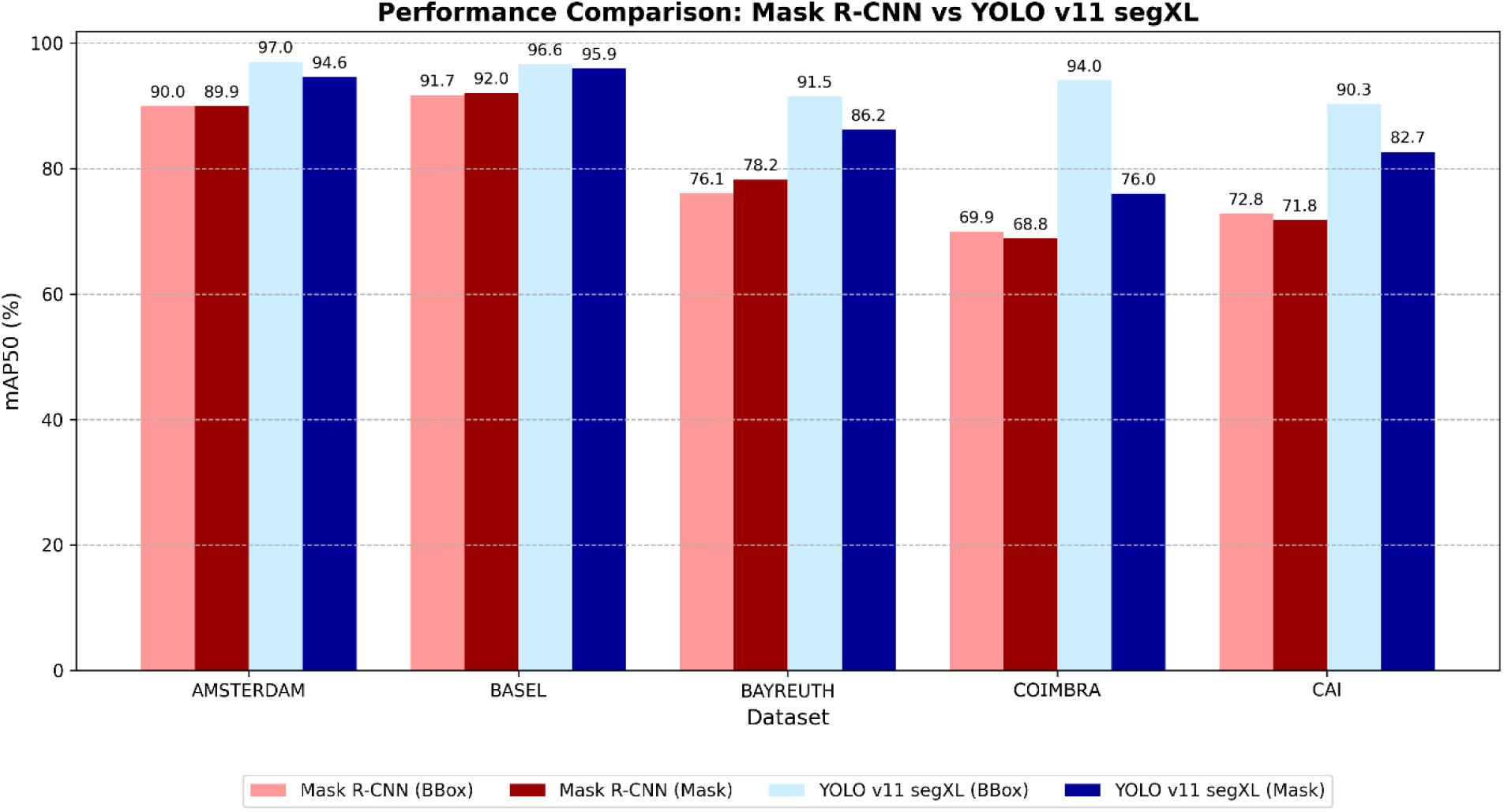
Comparative detection performance of YOLO v11 segXL and Mask R-CNN for automated Collembola detection in ecotoxicological toxicity tests. Performance is evaluated across five datasets (Amsterdam, Basel, Bayreuth, Coimbra and CollembolAI representing different imaging setups and substrate conditions. Bars show mean average precision at 50% IoU (mAP50) for bounding box detection (BBox) and instance segmentation (Mask), highlighting differences in model accuracy under varying image quality and experimental conditions.

Under optimal imaging conditions, represented by the Basel dataset (Nikon D800, controlled lighting, high resolution), both models achieved high mAP50 values. The difference in BBox mAP50 was relatively small (YOLO 96.6 % vs Mask R-CNN 91.7 %), and Mask mAP50 was similarly close (95.9 % vs 92.0 %).

However, despite these comparatively modest mAP50 gaps, the F1-scores already revealed a substantial divergence: YOLO v11 segXL maintained a much better balance between precision and recall, whereas Mask R-CNN showed reduced precision, resulting in markedly lower F1 values. This indicates that even when localisation accuracy wass high for both models, Mask R-CNN produces more false positives.

A similar pattern was observed for the Amsterdam dataset. While BBox mAP50 values were again relatively close at a high level (YOLO 97.0 % vs Mask R-CNN 90.0 %), F1-scores diverged more strongly, reflecting the same imbalance in Mask R-CNN predictions. In contrast, Mask mAP50 differences were moderate, consistent with both models being able to segment well under reasonably controlled conditions.

For challenging imaging conditions, differences in mAP50 became substantially larger and aligned closely with the F1 results. In the Bayreuth dataset (high ISO, noise), YOLO v11 segXL clearly outperformed Mask R-CNN in both BBox and Mask mAP50 (BBox 91.5 % vs 76.1 %; Mask 86.2 % vs 78.2 %). These large mAP50 gaps coincide with dramatic drops in F1 for Mask R-CNN, indicating that its performance degradation is not limited to localisation accuracy but reflects a general loss of robustness under noisy conditions.

The effect is even more pronounced for the Coimbra dataset, characterised by low resolution and small sample size. Here, the difference in BBox mAP50 is extreme (YOLO 94.0 % vs Mask R-CNN 69.9 %), with Mask mAP50 also clearly lower. These large mAP50 discrepancies are mirrored by very low F1-scores for Mask R-CNN, driven by extremely low precision despite high recall. This demonstrates that Mask R-CNN struggles fundamentally with low-resolution soil ecotoxicological images, rather than merely failing at strict IoU thresholds.

Finally, the CollembolAI dataset shows an intermediate case. While mAP50 differences remained substantial (BBox 90.3 % vs 72.8 %; Mask 82.7 % vs 71.8 %), they are smaller than in Coimbra and Bayreuth. Correspondingly, F1-scores indicate reduced but still meaningful performance gaps, suggesting that the synthetic multiscale data partially mitigate extreme failure modes while still favoring YOLO v11 segXL.

Overall, mAP50 and F1 convey complementary but consistent information. In high-quality datasets, mAP50 values can appear relatively close between models, whereas F1-scores reveal meaningful differences in prediction reliability. In lower-quality or more challenging datasets, both mAP50 and F1 diverge strongly, clearly identifying YOLO v11 segXL as the more robust and reliable architecture.

Together, these metrics justify the selection of YOLO v11 segXL as the primary model for automated Collembola detection across heterogeneous ecotoxicological imaging conditions.

### Multiscale Training

To quantify the effect of multiscale training, we compared YOLO v11 segXL models trained with and without the Flatbug Collembola AI image pyramid (Figure 4). Performance was evaluated per dataset for both bounding boxes (BBox) and instance masks (Mask) using F1-score and mAP at IoU 0.5 and 0.5-0.95.

**Figure 4.**
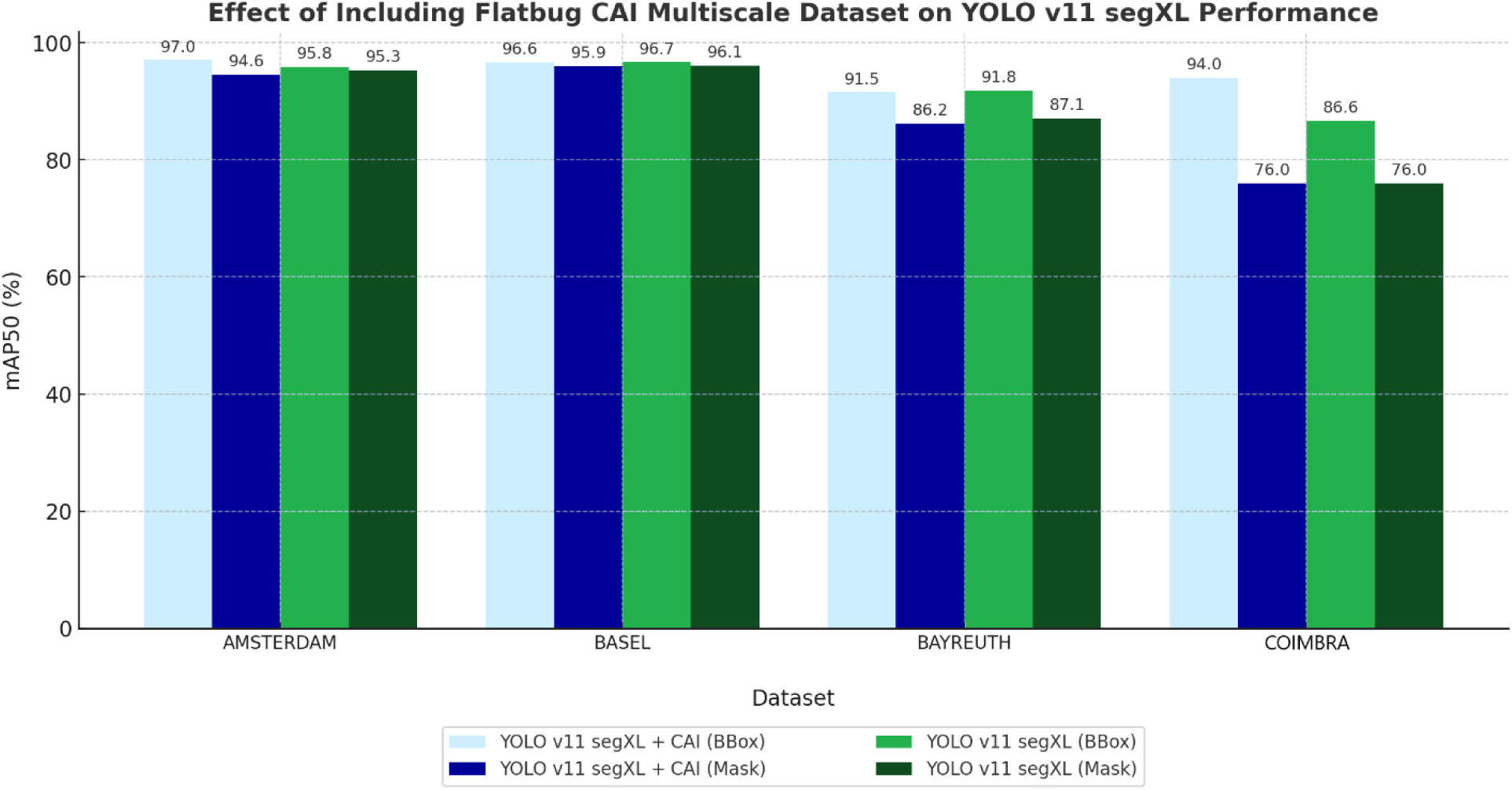
Effect of multiscale training with the CollembolAI dataset on YOLO v11 segXL performance for automated Collembola detection in eco-toxicological toxicity tests. Bars show mean average precision at 50% IoU (mAP50) for bounding box detection (BBox) and instance segmentation (Mask) across four datasets (Amsterdam, Basel, Bayreuth and Coimbra). Including the CollembolAI dataset during training improved detection accuracy under diverse imaging conditions, particularly for segmentation tasks.

Across the four high-resolution camera datasets, the inclusion of the CollembolAI pyramid produced small but consistent gains in BBox F1 on three of the four datasets, with slight decreases in Mask F1 (See Supplementary, Table 2):

Amsterdam (moderate quality): BBox F1 increased from 93.8% to 94.3% (+0.5 percentage points) and BBox mAP50 improved from 95.8% to 97.0%. In contrast, Mask F1 decreased slightly from 92.7% to 91.9%, and Mask mAP50 dropped from 95.3% to 94.6%.

Basel (highest image quality): Performance remained almost unchanged. BBox F1 slightly decreased from 94.6% to 93.8% (−0.8 pp), while Mask F1 dropped from 93.9% to 93.3% (−0.6 pp). BBox mAP50 was essentially identical (96.7% without vs. 96.6% with CollembolAI) and Mask mAP50 decreased only marginally (96.1% to 95.9%). At the stricter multiIoU criterion (mAP50-95), both BBox and Mask metrics showed small positive changes (+0.4 and +0.2 pp, respectively), indicating that the geometric quality of detections was at least preserved.

Bayreuth (high ISO, noisy): BBox F1 increased from 86.7% to 87.1%, reflecting a small net improvement. This corresponds to a modest precision-recall tradeoff: precision decreased (from 92.0% to 88.9%), while recall increased (from 81.9% to 85.4%), indicating that the CollembolAI-augmented model detected more true positives at the cost of slightly more false positives. Mask F1 decreased from 83.7% to 82.7% and Mask mAP50 dropped from 87.1% to 86.2%, suggesting a minor reduction in segmentation robustness for this challenging dataset.

Coimbra (low resolution, small sample size): The strongest positive effect of multiscale CollembolAI training was observed for the Coimbra dataset. BBox F1 increased from 84.6% to 85.8%, and BBox mAP50 improved markedly, from 86.6% to 94.0% (+7.4 pp). BBox mAP50-95 also increased from 41.4% to 44.7% (+3.3 pp), indicating more stable localization performance across IoU thresholds. Mask F1, in contrast, decreased from 75.7% to 72.4% (−3.2 pp), with little change in Mask mAP50 (∼76.0% for both models). This is consistent with a strong increase in Mask precision (from 73.1% to 80.2%) but a substantial drop in recall (from 78.5% to 66.0%), i.e. fewer spurious masks but more missed individuals. CollembolAI test set showed high performance under the CollembolAI-augmented model, with a BBox F1 of 88.0%, BBox mAP50 of 90.3%, and BBox mAP50-95 of 74.7%. Mask performance on CollembolAI also remained strong (Mask F1 80.5%, mAP50 82.7%, mAP50-95 56.8%), demonstrating good detection and segmentation on synthetic multiscale conditions with uniform backgrounds and high organism densities.

### Comparison Manual vs Automated

Across all datasets, the automated counts strongly correlated with manual counts, with R² values ranging from approximately 0.88 to 0.99, See also Supplementary Table S3. Additionally, the regressionlines were similar to the 1:1 lines (Automatic : Manual), See figure 5. This variation reflects dataset-specific factors such as image quality, sample preparation, and general image conditions. For example, datasets from Amsterdam, Basel and Denmark show high agreement (R² > 0.95), while a few datasets, such as Coimbra Boric Acid OECD 5, exhibit slightly lower R² values (∼0.88), due to differences in experimental setup or image characteristics.

**Figure 5.**
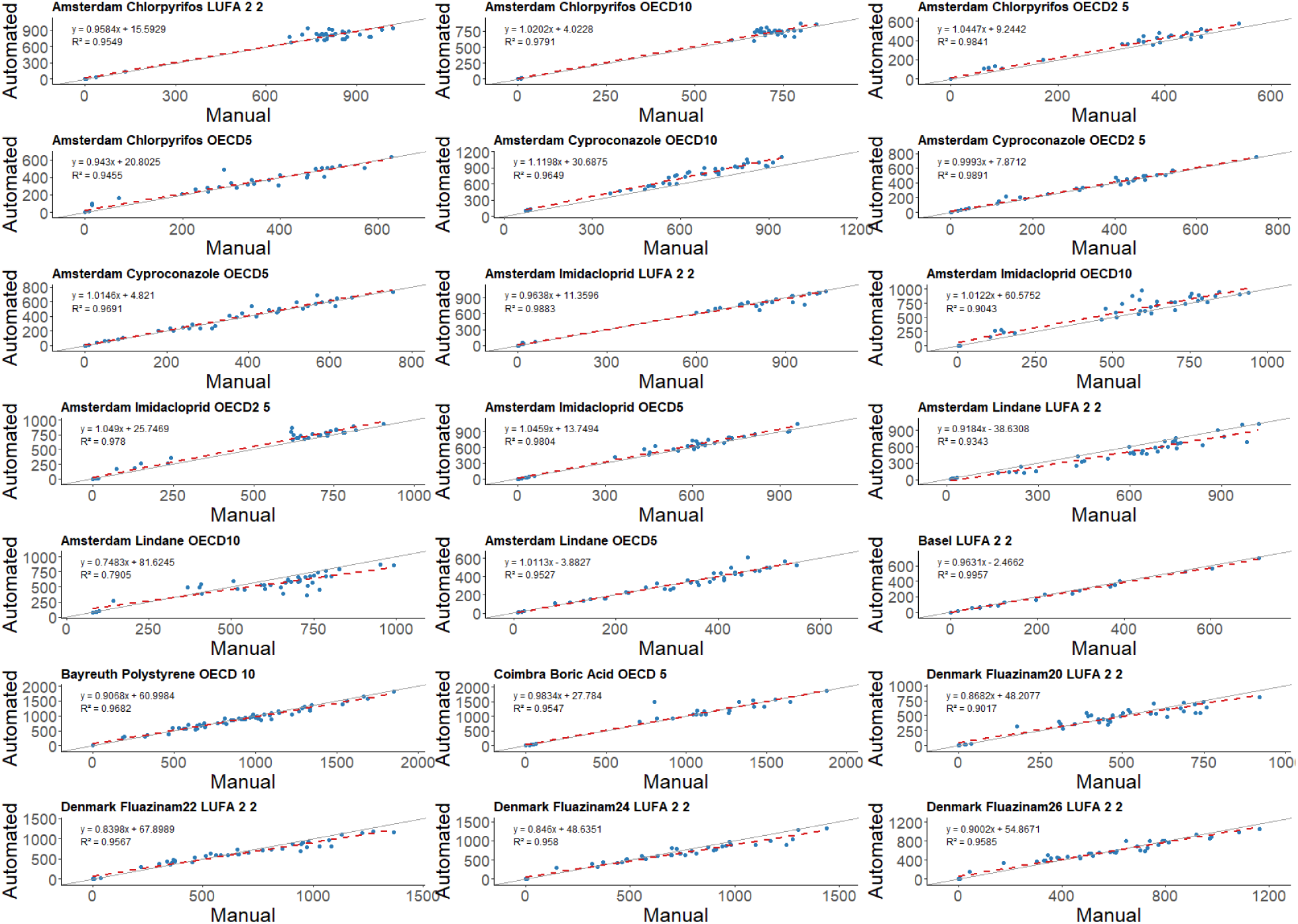
Correlation between automated and manual counts of collembolans across multiple datasets representing different laboratories and soil types. Each panel shows a linear regression fit (dashed red line) with the corresponding regression equation and coefficient of determination (R²), alongside the 1:1 identity line (solid line).

Dose–response curves (Figure 6) fitted to data derived from both counting methods strongly overlapped, capturing the expected decline in juvenile abundance with increasing concentrations. Differences in curve steepness or upper asymptotes were minimal and did not affect overall trend interpretation. Effect concentration estimates (EC_10_–EC_90_) were generally consistent between manual and automated approaches, with most differences falling within 95% confidence intervals. Notable exceptions included Amsterdam imidacloprid OECD10, where the automated approach resulted in a higher EC_50_ estimate and Amsterdam imidacloprid OECD2.5, which showed a slight shift. Conversely, Amsterdam Lindane LUFA 2.2 exhibited a shift toward a more conservative EC_50_ under the automated method.

**Figure 6.**
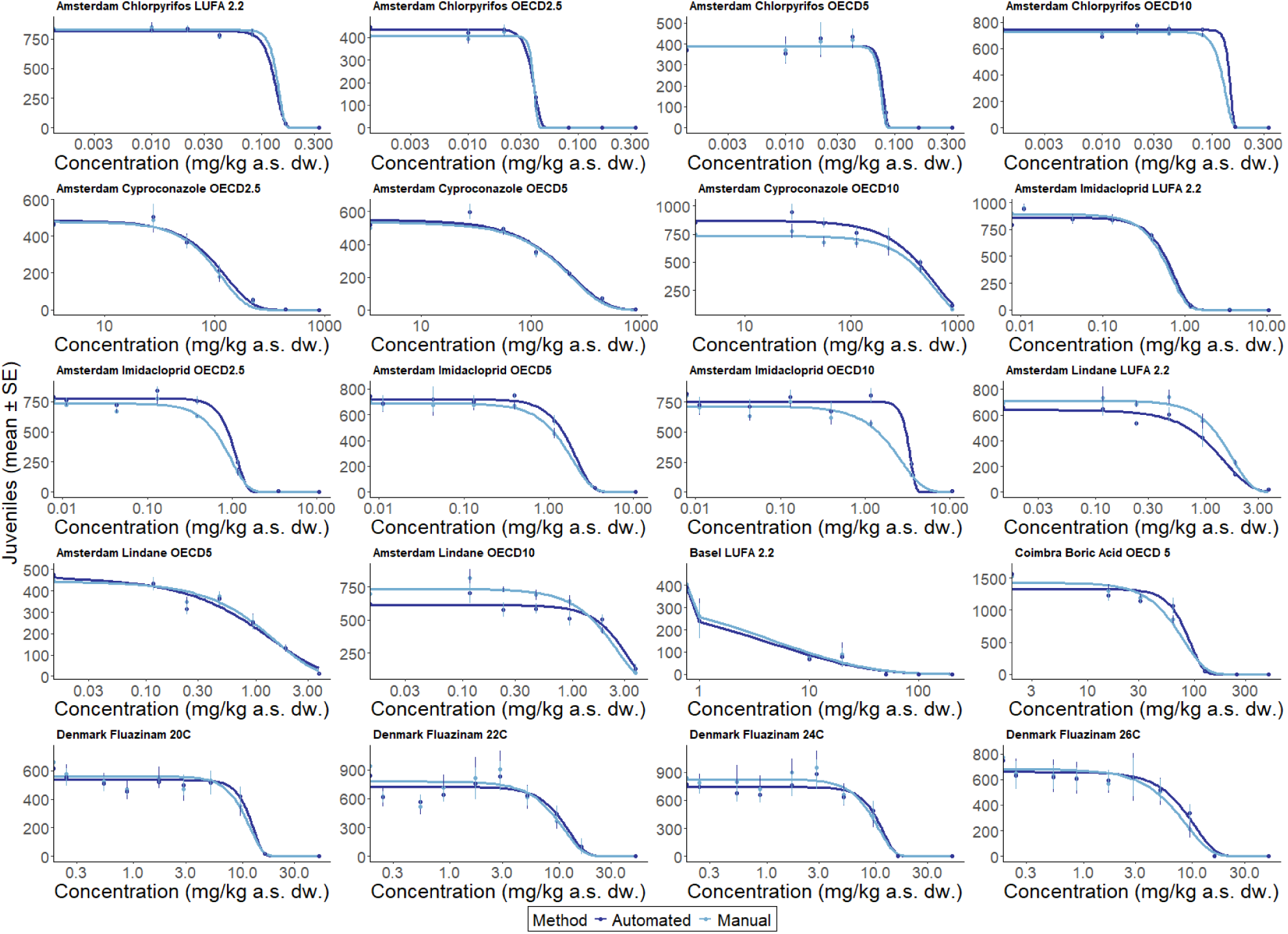
Comparison of dose–response curves for the effect of pesticides on springtail (*Folsomia candida*) reproduction obtained by using manual and automated juvenile counts across tested substances, fitted with three-parameter Weibull and LogLogistic models.

The comparison between manual and automated estimates (figure 7) of effective doses (EC_10_, EC_20_, EC_50_ and EC_90_) demonstrated an exceptionally strong agreement across all endpoints. Regression analyses revealed near-perfect linear relationships, with coefficients of determination (R²) exceeding 0.977 for all EC levels. The slopes of the fitted lines ranged from 0.8556 (EC_10_) to 1.0533 (EC_90_), indicating minimal deviation from the identity line (y = x). Intercepts were close to zero, further confirming consistency between the two methods. EC_x_ values differed minimally with a median %Δ of 6.2 ± 23 and overlap of automated and manually calculated EC_10_ -EC_90_ values of R^2^ ≥ 0.977. These differences correspond to a factor range of approximately 0.83 to 1.29 (median factor ≈ 1.06), indicating that automated estimates deviate by less than 30% from manual counts.

**Figure 7.**
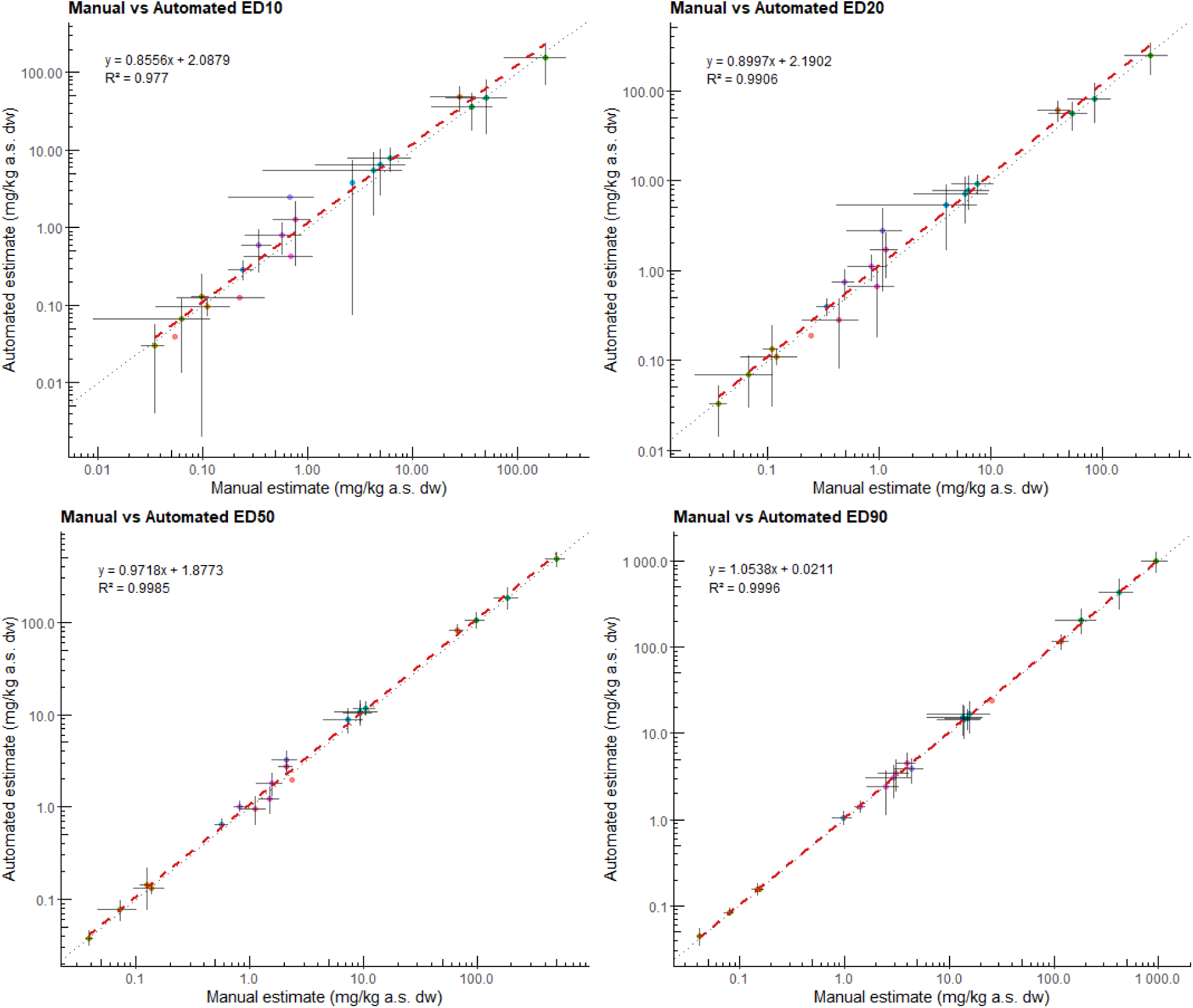
Comparison of effect concentration estimates (EC10–EC90) derived from manual and automated counts across substances. Each panel shows the relationship between manual and automated estimates for EC10, EC20, EC50 and EC90, including 95% confidence intervals. The dotted line represents the 1:1 identity line, while the dashed red line indicates the linear regression fit. Regression equations and R^2^ are depicted in the top left corner.

For lower effect levels (EC_10_ and EC_20_), automated estimates tended to be slightly higher than manual estimates as reflected by slopes below 1. Conversely, at EC_90_, the slope slightly exceeded 1, suggesting a minor tendency for automated estimates to arrive at higher juvenile numbers at higher doses at the upper tail of the dose-response curve. Confidence intervals (error bars) around the effect concentrations were generally overlapping between methods, although wider intervals were observed for EC_10_ and EC_20_, reflecting greater uncertainty at low effect levels.

The analysis of NOEC and LOEC values across substances revealed clear concentration-dependent effects on the reproduction of *Folsomia candida*. Statistical assumption checks determined whether parametric or non-parametric approaches were applied. NOEC and LOEC values (rounded to three decimal places) are summarized in Supplementary Table S5. Across substances, automated identification of NOEC and LOEC thresholds was consistent with manual assessments (Supplementary Figure S4). For most chemicals, the automated method produced identical thresholds to those determined by human counting. Some deviations occurred for the Denmark 20, 22 and Amsterdam Imidacloprid OECD 2.5 and 10 datasets, where automated identification resulted in higher NOECs (Denmark: 9.44 from 5.06 mg kg^-1^dry soil; Amsterdam: 1.169 from 0.011 and 0.390 mg kg^-1^ dry soil) compared to manual evaluation.

### Visual Performance

In the Amsterdam test dataset example (figure 8), the automatic detections aligned closely with the visible Collembola on the brown substrate, accurately capturing almost all individuals. Background distractions, such as debris, the glass rim and the label on the left side, were correctly ignored, demonstrating minimal false positives outside the intended test area. Foam or reflections on the rim can cause false positives in rare instances. In regions of particularly high density, such as the central cluster, some individuals were merged into larger grouped detections. The *Hough-circle* accurately identified the region of interest, closely following the dish’s inner rim.

**Figure 8.**
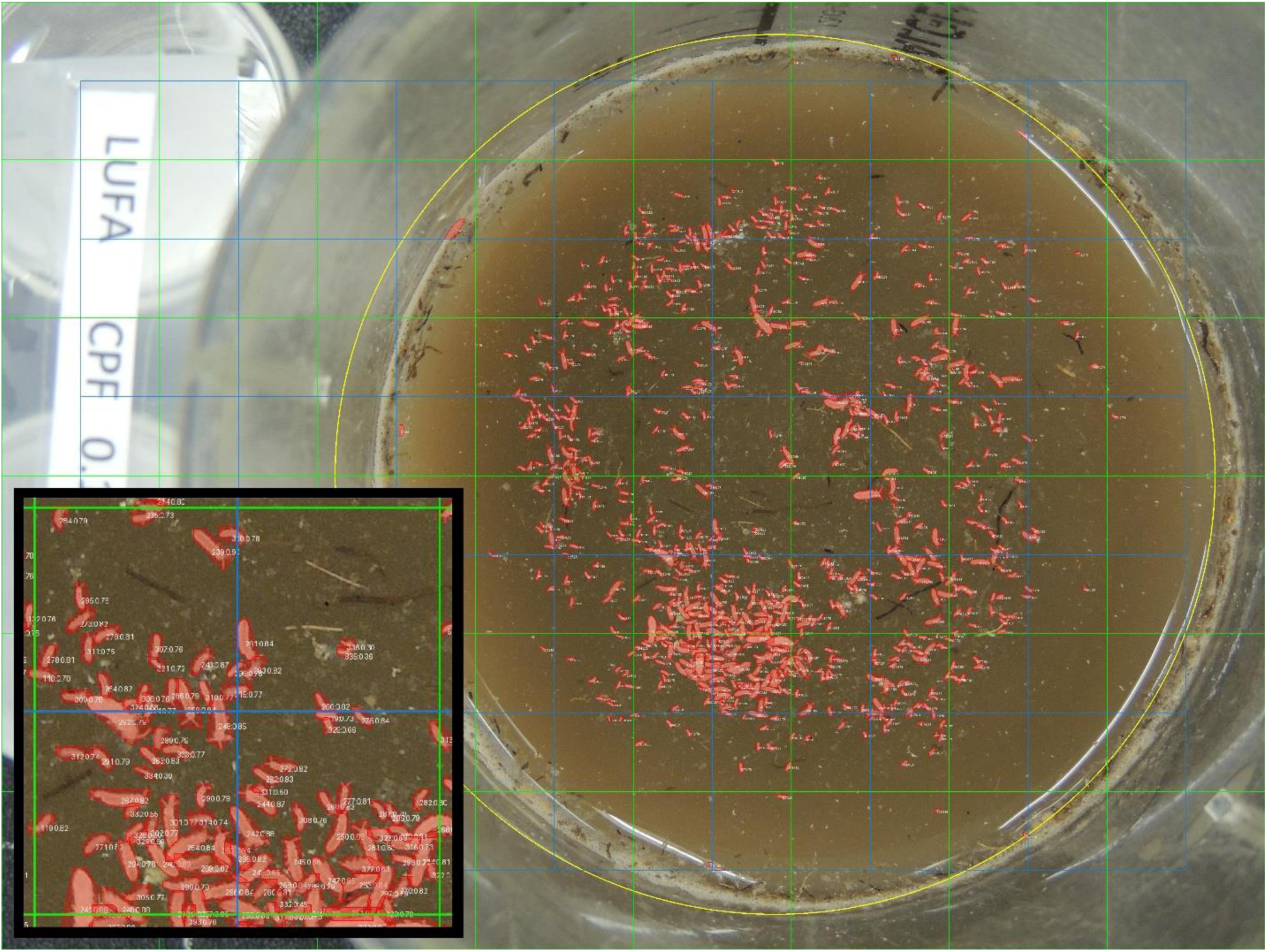
Detection performance example on Amsterdam dataset: accurate segmentation of Collembola on brown substrate with minimal false positives outside the region of interest.

In the pre-masked Bayreuth dataset test example (figure 9), the detection performance on a neutral, uniformly masked surrounding was highly accurate, consistently capturing individual Collembola without noticeable false positives. Even though the brighter fluid medium presented a lower contrast to the Collembola segmentation, performance was high. Regarding the *Hough-circle* region of interest detection, the circle was approximately centered but not precisely fitted. At the upper region, the circle slightly extended beyond the dish, including some empty background, whereas at the lower edge it partially cutted into the dish, potentially excluding valid detections at the lower boundary and thus resulting in a marginal underestimation of the Collembola count.

**Figure 9.**
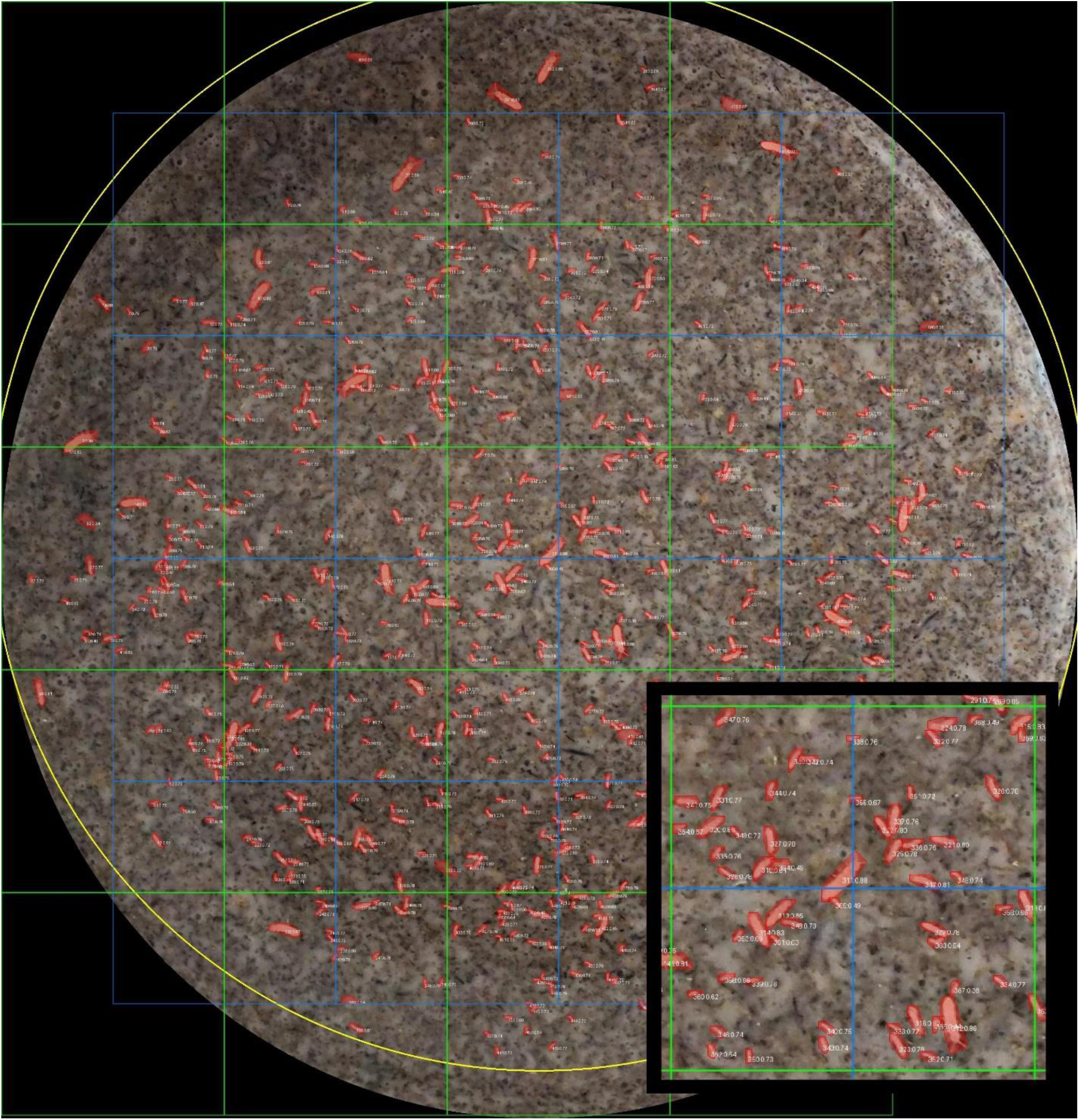
Detection performance example on Bayreuth dataset: accurate segmentation of Collembolans, despite low contrast and foam presence; slight region of interest circle misalignment noted.

The non-standardized Basel test dataset example (figure 10) demonstrated strong detection performance within the test jar on the dark substrate, successfully identifying most Collembola individuals. Few false positives detections appeared in areas with reflections or caustics. With an off-center positioning of the dish and a heterogenous background, the *Hough-circle* method failed to delineate a circle in this non-standardized setup. Consequently, the area of interest was not constrained, and this scenario suffered from noticeable false positives due to detections incorrectly marking printed text labels situated outside the test jar as well as the glass rim and some yellow duct tape (marked in red color), demonstrating a sensitivity to external interference.

**Figure 10.**
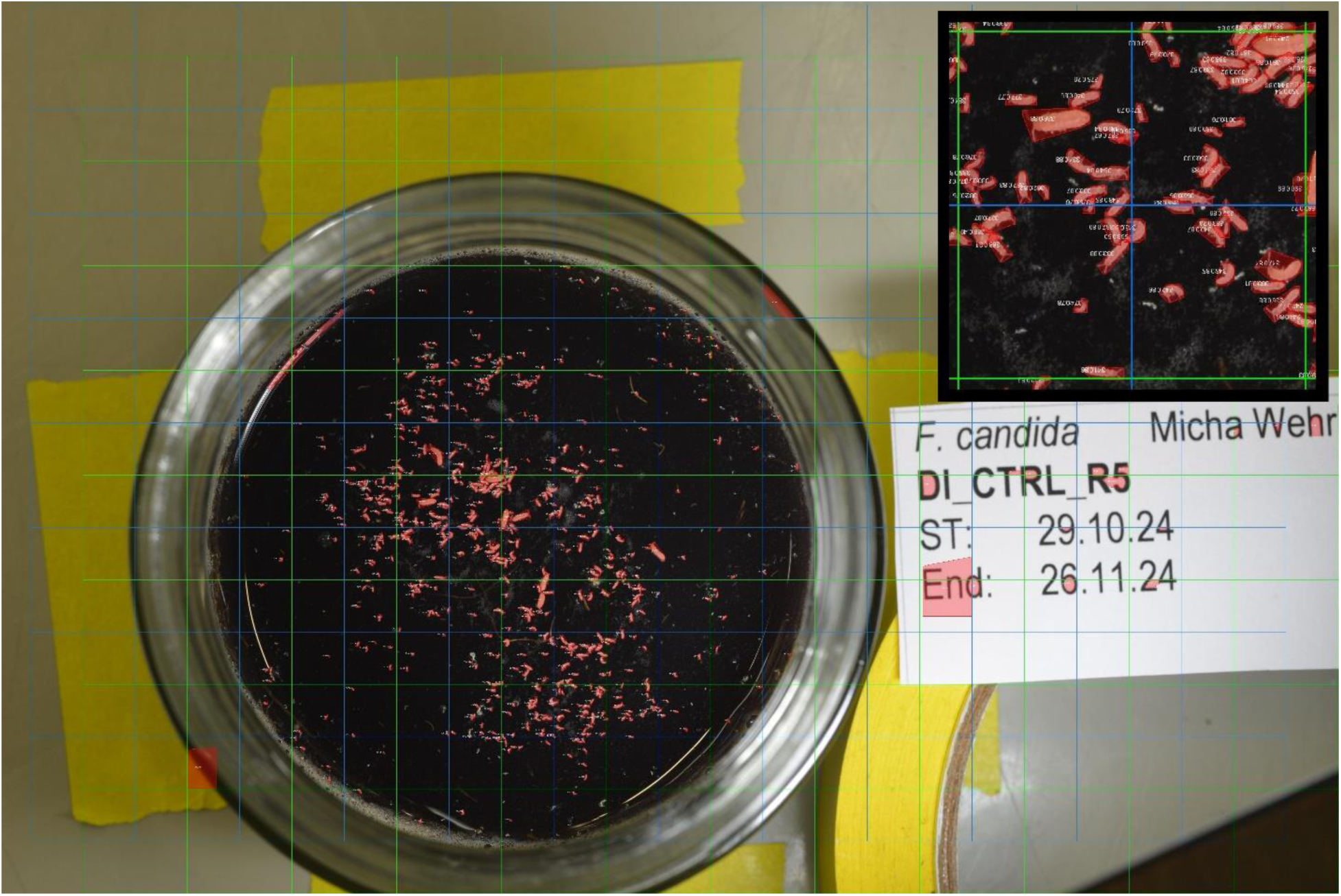
Detection performance example on non-standard Basel dataset: strong detection of Collembolans within jar but increased false positives due to off-center dish and external labels with letters.

Despite being slightly out of focus, this standardized Basel dataset test example (figure 11) exhibited a high detection performance on the dark black-brown substrate. Collembola individuals were accurately detected and outlined, even in dense clusters, reflecting excellent detection precision. Importantly, no external elements, such as the ruler or the surrounding table surface, triggered false positives, emphasizing high specificity. The *Houghcircle* region of interest detection here produced a circle closely fitting the inner edge of the dish and effectively encompassing the entire relevant soil surface.

**Figure 11.**
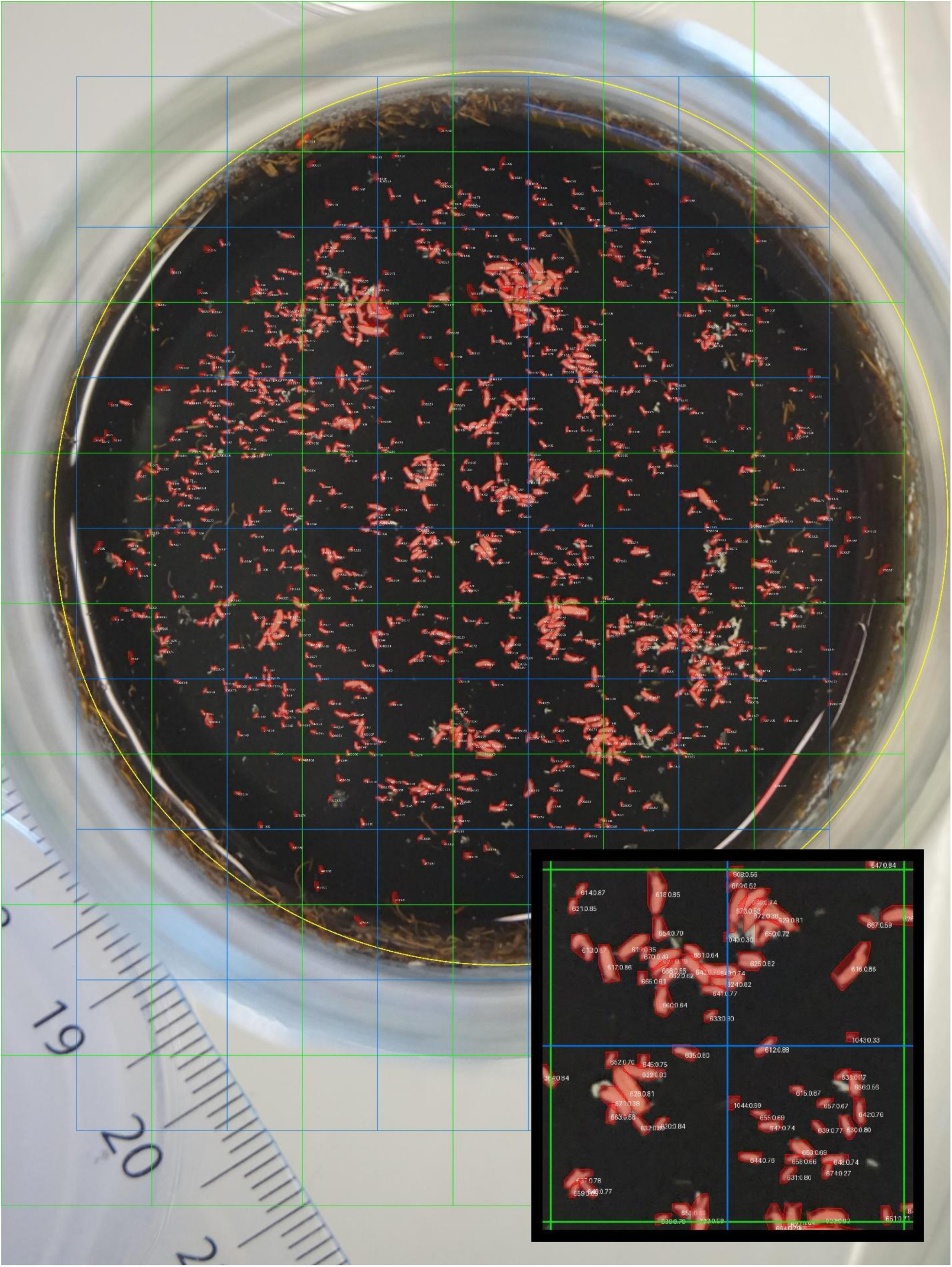
Detection performance example on standardized Basel dataset: high precision and specificity even in dense clusters of Collembolans; region of interest circle accurately fitted.

## Discussion

### General discussion of model performance

To evaluate the impact of multiscale augmentation, we compared two YOLO v11–xl–seg models: (i) a baseline model trained only on high-resolution OECD camera datasets, and (ii) a CAI-augmented model trained with additional multiscale tiles from the CollembolAI dataset (Sys et al., 2022). This comparison tested whether exposure to systematic variation in object scale and density improved robustness under diverse imaging conditions. We tried to also include an additional dataset from flatbug (Svenning et al., 2025), but the model performed significantly worse for our intent, thus we decided to only include the CollembolAI subset (Sys et al., 2022). Higher pyramid levels (L2-L16) produce image tiles with smaller apparent body sizes and denser populations on a uniform background. Conceptually, this should encourage the model to learn scale robust features and remain reliable when collembolans appear at different magnifications, resolutions and densities than those present in the high-resolution camera datasets alone. The empirical results support this intuition primarily at the level of object detection (bounding boxes).

The strongest effect of the CollembolAI-augmented model showed the strongest improvement for the Coimbra dataset, which is characterized by lower resolution, motion blur and uneven illumination. Here, BBox mAP50 increased by 7.4 percentage points and BBox mAP50-95 by 3.3 points when CollembolAI was included. This indicates that the CollembolAI-augmented model localizes Collembola much more reliably in difficult, low-resolution conditions. For Amsterdam and Bayreuth data sets, BBox F1 increased by around 0.5 percentage points, suggesting a modest but consistent gain in detection accuracy, even when imaging conditions were only moderately or severely degraded.

In contrast to detection, Mask F1 slightly decreased for most datasets when CollembolAI was included (−0.6 to −3.2 pp). The effect was modest for high-quality images (Basel, Amsterdam) and more pronounced for images representing lower quality like those from Coimbra. The precision-recall patterns indicate that the segmentation head became more conservative: precision often increased (fewer false positive masks), while recall decreased (more missed individuals). This is plausible given the domain shift between CollembolAI (uniform dark background, high contrast) and the more complex, heterogeneous backgrounds in the camera datasets. The segmentation head appeared to adapt to the simpler CollembolAI boundary conditions, which slightly reduced its tendency to segment partially visible or low-contrast individuals in noisy conditions.

The Basel dataset represented the best imaging conditions among all datasets we analyzed. Its metrics changed minimally with the CollembolAI-augmented model and remained the highest overall for Mask metrics (Mask F1 ≈ 93-94%, Mask mAP50 ≈ 96%). This suggests that the model was already close to a performance ceiling under ideal conditions and that adding CollembolAI did not substantially degrade this upper bound. The small absolute changes were within the range typically attributable to annotation noise, sampling variation, or minor training stochasticity.

The baseline model without the CollembolAI images did not see any CollembolAI-like multiscale, high-density, uniform-background imagery during training. Even if some segmentation metrics on the highly standardized datasets were marginally lower, this added capability is crucial for processing existing and future lower resolution or quality imagery (e.g. for benchmarking or transferring models to other labs using similar scanning setups). It ensures robust performance in practical scenarios where magnification, scanning resolution, or organism density differ from the exact conditions represented by the Basel, Amsterdam, Bayreuth and Coimbra data sets.

For downstream density-response curve (DRC) analyses and automated counting, accurate detection and counting of individuals is typically more critical than perfectly delineated masks. The multiscale CollembolAI training systematically improved or maintained BBox performance on challenging datasets, particularly Coimbra, where robust counting from low-quality images was otherwise difficult. The slight reduction in mask quality did not materially affect the derived abundance estimates, while the improved detection stability across scales and camera systems directly benefitted ecological inference and cross-experiment comparability.

Given the intended use of the tool-robust, cross-platform automated counting and segmentation of collembolans across a wide range of imaging conditions, we opt for the model trained with the CollembolAI multiscale dataset as our main model. The CollembolAI-augmented model offers better generalization to variable scales and resolutions and supports a wider set of real-world applications, while retaining near optimal performance under ideal imaging conditions.

### Comparison to manual counting

Even when imaging conditions were not optimized, COLLEMBOT consistently produced reliable outputs, showing strong agreement with manual counts and effect metrics within acceptable and reliable confidence intervals for regulatory endpoints. Across all validation datasets, coefficients of determination (R²) between manual and automated counts typically exceeded 0.90 and effect concentration estimates (EC_10_–EC_90_) and NOEC/LOEC values derived from automated counts were highly comparable to manual assessments (EC_x_ Median %Δ 6.2 ± 23 and EC_10_ -EC_90_ overlap R^2^ ≥ 0.977, see figure 7). These findings confirm that COLLEMBOT can serve as a robust alternative to manual counting in OECD 232 tests without compromising scientific integrity.

Performance differences observed across datasets were primarily driven by image quality. The most influential factors include:

- Lighting and Contrast: Uniform illumination and minimal reflections significantly improve detection accuracy.
- Focus and Resolution: High-resolution, sharp images reduce false positives and improve segmentation, especially in dense clusters.
- Background Complexity: Foam, debris, or uneven coloration can obscure individuals and reduce contrast.
- Organism Density: Extremely high densities may lead to merged detections, while very low densities (including empty samples) can increase false positives.

To counteract these limiting factors, some simple, fast and cheap modifications for optimization can be done as described in the optimization chapter in Supplementary Information S2. When these practices are followed, COLLEMBOT delivers highly reliable outputs that are improved to non-optimized images. Under optimal imaging conditions, confidence in automated effect metrics is high, with deviations generally within the range of biological variability observed in manual assessments. Even under suboptimal conditions, performance remains robust, supporting its use in routine ecotoxicological assessments. Performance for suboptimal condition can also be enhanced with Real ESRGAN (GAN) (X. Wang et al., 2021) based AI uscaling, as described in Supplementary Information S3.

- High-quality images → High confidence (R² > 0.95; EC metrics within confidence intervals)
- Moderate image quality → Moderate confidence (R² ∼ 0.90; minor deviations at extremes)
- Poor image quality / foam → Low confidence (manual verification is recommended for at least 10% of the replicates to demonstrate that only minor deviations occur)

Further supported by reducing labor intensity by up to 97% and improving reproducibility, COLLEMBOT represents a transformative step toward scalable, standardized hazard data generation in soil ecotoxicology. Its compatibility with OECD workflows ensures practical applicability and supports broader adoption for GLP-compliant testing.

Automated counting delivers substantial efficiency gains. In our validation, a test comprising 300 images and up to 1,500 organisms was processed in less than three hours, compared to approximately ∼137 hours for manual counting, a reduction of about 97%. This time saving skilled personnel from repetitive tasks, reduces labor costs and enables resources to be redirected toward more complex analyses or additional testing, ultimately increasing hazard data generation. In addition, standardized automated counting minimizes interpersonal variability and eliminates fatigue-related errors that can occur during prolonged manual assessments. This ensures greater consistency and reliability of the data, improving comparability across studies and enhancing the robustness of ecotoxicological evaluations.

### Comparison to other programs already in use

Existing automated counting solutions for ecotoxicological tests often require specialized workflows or hardware setups that deviate from OECD guidelines. For example, some approaches rely on anesthesia, thermal imaging (Pang et al., 2023), or custom extraction protocols and contrast enhancing counting in ImageJ (Caridade et al., 2011; Krogh et al., 2008), which limit their applicability in standardized tests.

Others integrate with proprietary laboratory systems (Bánszegi et al., 2014), making them less accessible and less comparable for routine or small-scale studies. Additionally the currently available computer vision methods, are not suitable for OECD tests either due to them being made for another purpose (Oriol et al., 2024; Sys et al., 2022).

In contrast, COLLEMBOT is designed for seamless integration into current OECD 232 workflows. It operates on standard photographs taken at the end of the test without requiring additional steps or modifications. This compatibility ensures that laboratories can adopt automated counting without investing in new infrastructure or altering established protocols, thereby reducing barriers to implementation and promoting broader adoption.

### Limitations and challenges

Image quality emerges as a critical determinant of the model’s counting and segmentation performance, as clearly demonstrated across our evaluated datasets. High-quality images consistently yield accurate, reliable detections and precise segmentation results, as exemplified by the standardized Basel dataset, which was characterized by controlled photographic conditions including optimal camera settings, external flash lighting and a well-designed although cheap photography chamber. Conversely, image quality degradation, such as off-center dish placement, high ISO noise, uneven lighting conditions, motion blur, or focus plane misalignment markedly reduce model accuracy and stability. Specifically, foam present on the medium surface poses a challenge, obscuring or partially concealing Collembola and reducing contrast, which adversely impacts segmentation accuracy, even though detection performance remains relatively robust.

To ensure consistently high-quality images conducive to accurate automated Collembola counting, standardized imaging protocols are crucial. Optimized photographic conditions include the use of a high-resolution camera equipped with an external flash to provide balanced, shadow-free illumination, low ISO settings to reduce image noise and a narrower aperture to enhance depth of field and sharpness.

Furthermore, minimizing overhead lighting interference using dedicated photographic chambers significantly improves image uniformity and reduces distracting reflections, shadows or caustics. These practices not only increase the model’s precision and recall for both detection and segmentation but also reduce susceptibility to false positives arising from external background. Masking the area of interest can be beneficial but is not necessary given a high picture quality and adherence to recommendations.

Our detector currently exhibits the highest error rate when applied to blank controls and samples that do not contain visible Collembola or are low-quality images. Such cases frequently yield a significant number of false positives, as the system struggles to distinguish artifacts or background noise from genuine targets if none are present. Although we will release future versions with ongoing improvements focus on enhancing recall and reducing false positives rates, we currently advise users to preferentially apply our method to populated samples, while maintaining manual verification for trials known to be devoid of Collembola. Notably, this issue was absent in high-quality datasets, where lighting and focus were optimal.

### Further training possibilities for the COLLEMBOT model

The proposed model can be tailored to the target application environment by performing retraining with domain-specific labeled data using a semi-supervised labeling strategy. In this approach, custom images are incorporated into the processing pipeline, followed by automated segmentation. The segmentation results must subsequently be validated using an annotation tool such as “*LabelMe”*, ensuring that masks are meticulously checked and corrected for each individual sample. The finalized and corrected masks can then be integrated into the training dataset, enabling model retraining to enhance performance within the specified context.

Currently, the model detects absolute abundance but does not distinguish between juveniles and adults. Although the model cannot distinguish adults from juveniles, this has little impact on labor intensity, as counting the number of adults requires little effort. Future iterations could incorporate size-based classification, leveraging the significant size difference between adults (introduced at test start) and offspring after 28 days. Additionally, size estimation could assume a cylindrical body shape to calculate surface area and length, providing the option to gain valuable information on growth-related effects in toxicity studies (Bánszegi et al., 2014; Gruss et al., 2022, 2024). Furthermore, the current model was trained primarily on *Folsomia candida,* but can possibly be extended to species with other species, such as *Folsomia fimetaria* or *Sinella curviseta*. The training dataset also includes 11 other species from the CollembolAI (Sys et al., 2022) dataset, but they have not been validated in the OECD test set up, yet. In a near-future, the model could also be made multi-class to identify egg clusters, detect and measure antenna length, or count the number of molts in a test jar, as demonstrated for *Hyalella* individuals (Pineda-Alarcón et al., 2023). These enhancements would allow extraction of previously inaccessible data from OECD tests and other studies, significantly expanding the ecological insights obtainable from standard toxicity assays. The model updates will be noted on the COLLEMBOT GitHub repository.

### Potential for standardization acceptance potential

For automated counting to be accepted and integrated in an international standard, formal ring-testing across multiple laboratories is required. We have shown in this paper that COLLEMBOT works on all OECD-mentioned standard soils and for the specified species *Folsomia candida*, with tests conducted in five different laboratories. Because the model operates on images captured according to OECD 232 guidelines, such ring tests are feasible without modifying existing protocols. Furthermore, the availability of cloud-based platforms such as Huggingface eliminates the need for expensive local hardware, making the approach accessible for independent research and inter-laboratory validation.

In addition, for implementation in GLP-compliant tests, prior internal validation is required even if the method is not yet part of an international standard. This can be achieved by applying the automated counting method to replicates with pre-counted collembolans and comparing results to manual counts. Such validation ensures reliability and compliance, similar to the process already performed for manual counting in many facilities.

## Conclusion

COLLEMBOT offers an open-source, robust and scalable solution to one of the most labor-intensive steps in soil ecotoxicological testing: manual counting collembolans. Minor discrepancies observed under challenging imaging conditions are primarily attributable to image quality factors, such as contrast, foam and focus, rather than systematic model bias. By reducing analysis time by up to 97%, improving reproducibility and maintaining strong agreement with manual counts (R² > 0.88 - 0.99), COLLEMBOT addresses a major bottleneck in hazard data generation. Its compatibility with existing OECD protocols ensures seamless integration into current workflows without procedural changes.

We encourage researchers and regulatory bodies to adopt and further develop COLLEMBOT. The model is validated on all currently used standard soils and supports *Folsomia candida* mentioned in the OECD 232 guideline, offering a true one-to-one replacement for manual counting.

By embracing automated counting, the ecotoxicological community can accelerate data generation, reduce costs, and improve science-based decision-making for soil organism protection. Collaborative development will be key to expanding COLLEMBOT’s capabilities and ensuring its acceptance in GLP-compliant testing.

## Data Availability Statement

The model weights generated and analyzed during the current study, and the datasets and scripts for dose–response modeling is available in the Zenodo repository at DOI: https://doi.org/10.5281/zenodo.17987887. The code for the model training and inference is available at GitHub at https://github.com/waldstrom/collembot. All data not publicly available can be obtained from the corresponding author upon reasonable request.

## Acknowledgements

We strongly thank J.G. Honoré for the design of the graphical abstract.

## CRediT Author Contributions

Conceptualization: Micha Wehrli, Adrian Meyer, Miriam Langer

Methodology: Micha Wehrli, Adrian Meyer, Magdalena Mair

Software: Adrian Meyer

Validation: Micha Wehrli, Adrian Meyer, Magdalena Mair

Formal analysis: Micha Wehrli, Adrian Meyer

Investigation: Micha Wehrli, Adrian Meyer, Éverton Souza da Silva

Resources: All authors

Data curation: Micha Wehrli, Adrian Meyer, Éverton Souza da Silva

Writing – original draft: Micha Wehrli, Adrian Meyer, Éverton Souza da Silva, Miriam Langer

Writing – review & editing: All authors

Visualization: Adrian Meyer, Micha Wehrli

Supervision: Miriam Langer, Kees Van Gestel, Denis Jordan, Magdalena Mair

Project administration: Micha Wehrli, Miriam Langer

Funding acquisition: Miriam Langer

## References

1. Abreu, S. N., Jesus, F., Domingues, I., Baptista, F., Pereira, J. L., Serpa, D., Soares, A. M. V. M., Martins, R. E., & Oliveira E Silva, M. (2022). Automated Counting of Daphnid Neonates, *Artemia* Nauplii, and Zebrafish Eggs: A Proof of Concept. Environmental Toxicology and Chemistry, 41(6), 1451–1458. 10.1002/etc.5323

2. Akiba, T., Sano, S., Yanase, T., Ohta, T., & Koyama, M. (2019). Optuna: A Next-generation Hyperparameter Optimization Framework (No. arXiv:1907.10902). arXiv. 10.48550/arXiv.1907.10902

3. Bánszegi, O., Kosztolányi, A., Bakonyi, G., Szabó, B., & Dombos, M. (2014). New Method for Automatic Body Length Measurement of the Collembolan, Folsomia candida Willem 1902 (Insecta: Collembola). PLOS ONE, 9(6), e98230. 10.1371/journal.pone.0098230

4. Boykov, Y. Y., & Jolly, M.-P. (2001). Interactive graph cuts for optimal boundary & region segmentation of objects in N-D images. Proceedings Eighth IEEE International Conference on Computer Vision. ICCV 2001, *1*, 105–112 vol.1. 10.1109/ICCV.2001.937505

5. Caridade, C. M. R., Marcal, A. R. S., Mendonca, T., Natal-da-Luz, T., & Sousa, J. P. (2011). Automatic counting the number of Collembola in digital images. 2011 4th International Congress on Image and Signal Processing, 4, 1837–1841. 10.1109/CISP.2011.6100624

6. De Souza Machado, A. A., Kloas, W., Zarfl, C., Hempel, S., & Rillig, M. C. (2018). Microplastics as an emerging threat to terrestrial ecosystems. Global Change Biology, 24(4), 1405–1416. 10.1111/gcb.14020

7. EC. (2009). Regulation (EC) No 1107/2009.

8. Gruss, I., Lallaouna, R., Twardowski, J., Magiera-Dulewicz, J., & Twardowska, K. (2024). Collembola growth in heavy metal-contaminated soils. Scientific Reports, 14(1), 27998. 10.1038/s41598-024-79766-5

9. Gruss, I., Twardowski, J., Karczewska, A., Szopka, K., Kluczek, K., & Magiera-Dulewicz, J. (2022). Collembola reduce their body sizes under arsenic contamination in the soil – Possible use of new screening tool in ecotoxicology. Ecological Indicators, 142, 109185. 10.1016/j.ecolind.2022.109185

10. Hampton, L. M. T., Wyler, D. B., Almroth, B. C., Coffin, S., Cowger, W., Doyle, D., Hataley, E. K., Hutton, S. J., Mair, M. M., Miller, E. L., Monclús, L., Sharpe, E. E., Samreen, S., Ahmed, K. T., Allamby, Q. P. V., Vital, A. L. A., Asnicar, D., Bare, J. L., Barrick, A., … Mehinto, A. C. (2025). The Toxicity of Micro-plastics Explorer (ToMEx) 2.0. Microplastics and Nanoplastics, 5(1), 38. 10.1186/s43591-025-00145-6

11. He, K., Gkioxari, G., Dollár, P., & Girshick, R. (2018). *Mask R-CNN* (No. arXiv:1703.06870). arXiv. 10.48550/arXiv.1703.06870

12. Ioannou, D., Huda, W., & Laine, A. F. (1999). Circle recognition through a 2D Hough Transform and radius histogramming. Image and Vision Computing, 17(1), 15–26. 10.1016/S0262-8856(98)00090-0

13. Khanam, R., & Hussain, M. (2024). *YOLOv11: An Overview of the Key Architectural Enhancements* (No. arXiv:2410.17725). arXiv. 10.48550/arXiv.2410.17725

14. Krogh, P. H., João, M., Amorim, B., Andrés, P., Bakonyi, G., Becker Van Slooten, K., Domene, X., Geujin, I., Kaneko, N., Knäbe, S., Kocí, V., Lana, J., Moser, T., Princz, J., Schaefer, M., Scott-Fordsmand, J. J., Stubberud, H., & Wilke, B.-M. (2008). Toxicity testing with the collembolans Folsomia fimetaria and Folsomia candida and the results of a ringtest With contributions from.

15. Krogh, P. H., Johansen, K., & Holmstrup, M. (1998). Automatic counting of collembolans for laboratory experiments. Applied Soil Ecology, 7(2), 201–205. 10.1016/S0929-1393(97)00043-7

16. Lead, J. R., Batley, G. E., Alvarez, P. J. J., Croteau, M.-N., Handy, R. D., McLaughlin, M. J., Judy, J. D., & Schirmer, K. (2018). Nanomaterials in the environment: Behavior, fate, bioavailability, and ef-fects—An updated review. Environmental Toxicology and Chemistry, 37(8), 2029–2063. 10.1002/etc.4147

17. OECD. (2016a). OECD/OCDE 232 OECD GUIDELINES FOR TESTING CHEMICALS Collembolan Reproduction Test in Soil. http://www.oecd.org/termsandconditions/.

18. OECD. (2016b). Test No. 222: Earthworm Reproduction Test (Eisenia fetida/Eisenia andrei). OECD. 10.1787/9789264264496-en

19. OECD. (2016c). Test No. 226: Predatory mite (Hypoaspis (Geolaelaps) aculeifer) reproduction test in soil.

20. OECD. 10.1787/9789264264557-en

21. Oriol, T., Pasquet, J., & Cortet, J. (2024). Automatic identification of Collembola with deep learning techniques. Ecological Informatics, 81, 102606. 10.1016/j.ecoinf.2024.102606

22. Pang, A., Nicol, A. M., Rutter, A., & Zeeb, B. (2023). Improved methods for quantifying soil invertebrates during ecotoxicological tests: Chill comas and anesthetics. Heliyon, 9(1), e12850. 10.1016/j.heliyon.2023.e12850

23. Pineda-Alarcón, L., Zuluaga, M., Ruíz, S., Mc Cann, D. F., Vélez, F., Aguirre, N., Puerta, Y., & Cañón, J. (2023). Automated software for counting and measuring Hyalella genus using artificial intelligence. Environmental Science and Pollution Research, 30(59), 123603–123615. 10.1007/s11356-023-30835-8

24. Posit team. (2025). RStudio: Integrated Development Environment for R. Posit Software, PBC, Boston, MA. URL http://www.posit.co/.

25. Redmon, J., Divvala, S., Girshick, R., & Farhadi, A. (2016). You Only Look Once: Unified, Real-Time Object Detection. 2016 IEEE Conference on Computer Vision and Pattern Recognition (CVPR), 779–788. 10.1109/CVPR.2016.91

26. Ritz, C., Baty, F., Streibig, J. C., & Gerhard, D. (2015). Dose-Response Analysis Using R. PLOS ONE, 10(12), e0146021. 10.1371/journal.pone.0146021

27. Russell, B. C., Torralba, A., Murphy, K. P., & Freeman, W. T. (2008). LabelMe: A Database and Web-Based Tool for Image Annotation. International Journal of Computer Vision, 77(1), 157–173. 10.1007/s11263-007-0090-8

28. Schneider, C. A., Rasband, W. S., & Eliceiri, K. W. (2012). NIH Image to ImageJ: 25 years of image analysis. Nature Methods, 9(7), Article 7. 10.1038/nmeth.2089

29. Svenning, A., Mougeot, G., Alison, J., Chevalier, D., Molina, N. C., Ong, S.-Q., Bjerge, K., Carrillo, J., Høye, T. T., & Geissmann, Q. (2025). *A General Method for Detection and Segmentation of Terrestrial Arthropods in Images* (p. 2025.04.08.647223). bioRxiv. 10.1101/2025.04.08.647223

30. Sys, S., Weißbach, S., Jakob, L., Gerber, S., & Schneider, C. (2022). CollembolAI, a macrophotography and computer vision workflow to digitize and characterize samples of soil invertebrate communities preserved in fluid. Methods in Ecology and Evolution, 13(12), 2729–2742. 10.1111/2041-210X.14001

31. van Hall, B. G., Sweeney, C. J., Bottoms, M., & van Gestel, C. A. M. (2025). The influence of soil organic matter content on the toxicity of pesticides to the springtail Folsomia candida. *Environmental Toxicology and Chemistry*, vgae048. 10.1093/etojnl/vgae048

32. van Hall, B. G., & van Gestel, C. A. M. (2025). Automated quantification of *Enchytraeus crypticus* juveniles in different soil types using RootPainter. Ecotoxicology and Environmental Safety, 289, 117482. 10.1016/j.ecoenv.2024.117482

33. van Loon, S., Xie, G., Svendsen, C., Kraak, M. H. S., de Jeu, L., Schut, N. C., Sprokkereef, E., Hurley, R., van Wezel, A. P., & van Gestel, C. A. M. (2025). Microplastics and PFAS as ubiquitous pollutants affect potencies of highly toxic chemicals in mixtures. Journal of Hazardous Materials, 500, 140493. 10.1016/j.jhazmat.2025.140493

34. Wang, X., Xie, L., Dong, C., & Shan, Y. (2021). *Real-ESRGAN: Training Real-World Blind Super-Resolution with Pure Synthetic Data* (No. arXiv:2107.10833). arXiv. 10.48550/arXiv.2107.10833

35. Wang, Z., Walker, G. W., Muir, D. C. G., & Nagatani-Yoshida, K. (2020). Toward a Global Understanding of Chemical Pollution: A First Comprehensive Analysis of National and Regional Chemical Inventories. Environmental Science & Technology, 54(5), 2575–2584. 10.1021/acs.est.9b06379

36. Wehrli, M., Slotsbo, S., Fomsgaard, I. S., Laursen, B. B., Gröning, J., Liess, M., & Holmstrup, M. (2024). A Dirt(y) World in a Changing Climate: Importance of Heat Stress in the Risk Assessment of Pesticides for Soil Arthropods. Global Change Biology, 30(10), e17542. 10.1111/gcb.17542

37. Wickham, H. (with Sievert, C.). (2016). ggplot2: Elegant graphics for data analysis (Second edition). Springer international publishing.

38. Xue, F., Zhang, T., Jia, Y., Zhao, X., Wu, L., Yin, D., Schiwy, A., & Hollert, H. (2025). *Developing a Deep Learning-Based Image Analysis Model for High-Throughput Micronucleus Assays: Genotoxicity as a Sediment Quality Indicator in East Taihu and Yangcheng Lakes, China* (SSRN Scholarly Paper No. 5460402). Social Science Research Network. 10.2139/ssrn.5460402

