## Supplementary for "COLLEMBOT: AI-Based Counting of Collembola for OECD 232 Tests"

*Contents of Tables:*

*Supplementary Table 1. Overview of all experiments used for COLLEMBOT development and validation* 8
*Supplementary Table 2. YOLO v11 segXL performance metrics (F1-scores) for bounding boxes and masks across datasets as depicted in figure 4, with and without Flatbug CAI augmentation.* 9
*Supplementary Table 3. Linear regression models comparing automated counts against manual counts of OECD 232 tests with Folsomia candida, for each dataset, including regression formula and coefficient of determination ($R^2$).* 9
*Supplementary Table 4. Effect concentration estimates (EC10–EC90) for manual and automated counts of Collembola across substances, including confidence intervals.* 10
*Supplementary Table 5. Summary of NOEC and LOEC values (mg kg-1 dry soil) for all tested substances and counting methods (Manual vs Automated), including the statistical approach applied (ANOVA + Dunnett or Kruskal–Wallis + Dunn). NOEC represents the highest concentration without a statistically*

*significant effect compared to the control and LOEC represents the lowest concentration with a significant effect (α = 0.05). All concentrations are rounded to three decimal places.* 17

*Contents of Figures:*

*Supplementary Figure 1. Photography chamber for standardized imaging of OECD 232 test jars, constructed from a white bucket for light diffusion and equipped with a fixed-height tripod and ruler for scale.* 5

*Supplementary Figure 2: The original panel on the left depicts a picture from the Coimbra dataset before the AI upscalind, while the GAN 2x picture on the right was after an AI upscaling (Wang et al., 2021). The AI upscaling was done to improve the quality of the picture to better detect springtails in low quality settings.* 7

*Supplementary Figure 3: Boxplot for the comparison of three different counting methods for the Coimbra dataset: Hand counted pictures serve as ground truth, Original Images counted with Collembot and GAN 2x upscaled images before counting with Collembot* 7

*Supplementary Figure 4: Comparison of NOEC and LOEC values for the toxicity of pesticides to Folsomia candida, obtained from automated versus manual counts. The dotted line represents the 1:1 identity line, while the solid orange line shows the log10-scale east squares regression fit between the two methods. Regression equations and R2 values are displayed for each panel.* 16

### Supplementary Information S1: Test specifications for unpublished test datasets

#### Test Organisms for unpublished test sets

Test organisms, *Folsomia candida* “Berlin Strain”, were obtained from a mass culture maintained at FHNW Muttenz, which originated from the Department of Ecoscience in Aarhus, which were used for the Aarhus, Denmark dataset (Wehrli et al., 2024). They have been cultured in a laboratory for more than 30 years (Simonsen & Christensen, 2001). The cultures were maintained at 20±0.1˚C (16/8 h light/dark cycle and 100% humidity), on a petri dish with plaster of Paris and active charcoal (8:1 w/w) and fed weekly ad libitum with baker's yeast.

*Folsomia candida* used for tests for the Bayreuth dataset were kindly provided by the group of Liliane Ruess (Ecology Group, HU Berlin) and kept in the laboratory in Bayreuth for 12 months. The cultures were maintained at 20˚C (12/12 h light/dark cycle and 70% humidity), on a petri dish with plaster of Paris and active charcoal (8:1 w/w) and fed weekly ad libitum with baker's yeast.

The Coimbra dataset was provided by Cloverstrategy Lda (a GLP-compliant company for laboratory ecotoxicological testing, Coimbra, Portugal) using *Folsomia candida* originally obtained approximately five years ago from laboratory cultures at Coimbra University (Portugal). The organisms were reared at 20±2°C under a 16:8 h light:dark photoperiod in plastic containers with the bottom filled with a mixture of plaster of Paris and activated charcoal at a ratio of 11:1 (w:w).

##### OECD 232 Test

The tests were performed in accordance with OECD Guideline 232. Briefly, 10 synchronized (10-12 days) *Folsomia candida* individuals were added to each test jar containing soil spiked with the desired test substance and concentration using an aspirator. Animals were fed *ad libitum* at the start of the test with baker’s yeast and incubated into a climate chamber with a 16:8h light cycle and 100% humidity. Test jars were aerated briefly (10-30 seconds) 3 times a week, weighed every 7 days to account for water loss, and soil moisture content was corrected by adding demineralized water if necessary. No water loss was observed during the experiment. One replicate, without springtails, per tested concentration was added to assess moisture content, pH, and soil concentrations of the used test contaminant at the start and end of the test. After 28 days, the organisms were floated by adding water to the test jars and a few drops of black or blue ink (Pelikan 4001 Brilliant Black or Parker Quink Blue) to color the background to enhance the contrast and facilitate counting. The test jars were then stirred gently for 30 seconds and subsequently left for 5 minutes, allowing the springtails to float to the surface. After the initial stirring, the test jars were stirred a second time and then allowed to settle to ensure a calm surface before being transferred to the photography chamber.

##### High Resolution Datasets

Our training and evaluation pipeline utilized four primary high resolution imagery datasets (see example images in Figure 1):

"Basel", consisting of 417 tiles from a Nikon D800 camera (6316×4912 px, F/8, ISO200, flash), see setup in Supplementary Information S1: Images generally exhibited the highest quality, in comparison to the other datasets, characterized by clear focus, minimal noise, precise object outlines and balanced illumination, benefiting from optimal photographic settings including controlled lighting (flash), low ISO and narrow aperture. Except for edge cases with foam, the background medium took a very dark black-brown color boasting strong contrast to the Collembola bodies. Because of the high image resolution of approximately 31 Mpx, adult Collembola covered larger areas of an image tile while juveniles were easily discernible.

"Amsterdam", comprising 91 tiles from a Nikon Coolpix P510 (4608×3456 px, F/3.9, ISO400, no flash): The images had moderate to good quality, showing adequate detail and clarity. Despite acceptable lighting conditions, the higher ISO and slower shutter speed without flash slightly affected sharpness and contrast. The background medium took a medium brown color offering a very monotonous contrast to the Collembola segments. Edge cases included bright string-like reflections of laboratory ceiling lights with caustic refractions (e.g. Example 5 in Figure 1).

"Bayreuth", with 192 tiles captured by an Olympus E-M10MarkII under challenging low-light conditions (4608×3456 px, F/3.5, ISO5000, no flash): Images showed notably lower quality due to high ISO settings, resulting in pronounced noise and reduced sharpness. Due to the development of a foam cover on the surface, the background medium often reduced the contrast to individual collembola segments (e.g. Example 1 in Figure 1).

"Coimbra", consisting of 27 tiles from a Panasonic Lumix CD-TZ90 (3840x2160 px, F/3.3-6.4, ISO80-3200, no flash): Images exhibited moderate to low image quality, impacted by limited resolution, motion blur, uneven lighting conditions and reduced contrast. Background medium consisted of patchy bright brown hues interspersed with blue ink stains (e.g. Example 3). Individual Collembola had very few pixels covered. Some pictures showed motion or partial out-of-focus blur artefacts. Ceiling light reflections and caustic refractions were also apparent (e.g. Example 2 in Figure 1).

### Supplementary Information S2: Optimization suggestions for best practice

In addition, to better facilitate the generation of quality images that allow for robust and reliable collembolan counts, we developed and present a low-cost photography chamber that is both reliable and easy to set up, which was used to photograph one part of this dataset. This photography chamber was designed to address the need to eliminate reflections and enable a smooth picture-taking process.

#### Camera Choice

Optimal detection accuracy depends on the quality of the pictures, which is partially determined by hardware used. Generally, a DSLR camera with at least an APS-C sensor, with a ring light or indirect illumination, with a limited exposure time of <1/100s to minimize motion blur is recommended. We recommend adjusting the camera's white balance to match the actual colors of the pictures to reality. It is also recommended to use low ISO settings to suppress reflections from overhead lighting. Pictures should be saved in a JPEG format. In our case the optimal results were obtained with a Nikon D800, with an AF-S Nikkor 60mm f/2.8 objective and a Nikon Speedlight SB-600 flash, in a rudimentary photography chamber described below.

#### Photography chamber

The chamber (figure 1) can be constructed from simple materials. A white bucket serves as the main component, diffusing flash light evenly across the water surface to eliminate reflections. A fixed-height tripod ensures stability and a consistent distance for size estimation. Inside the bucket, a rig can be glued to hold the test jar in place and an adjustable ruler should be added for scale.

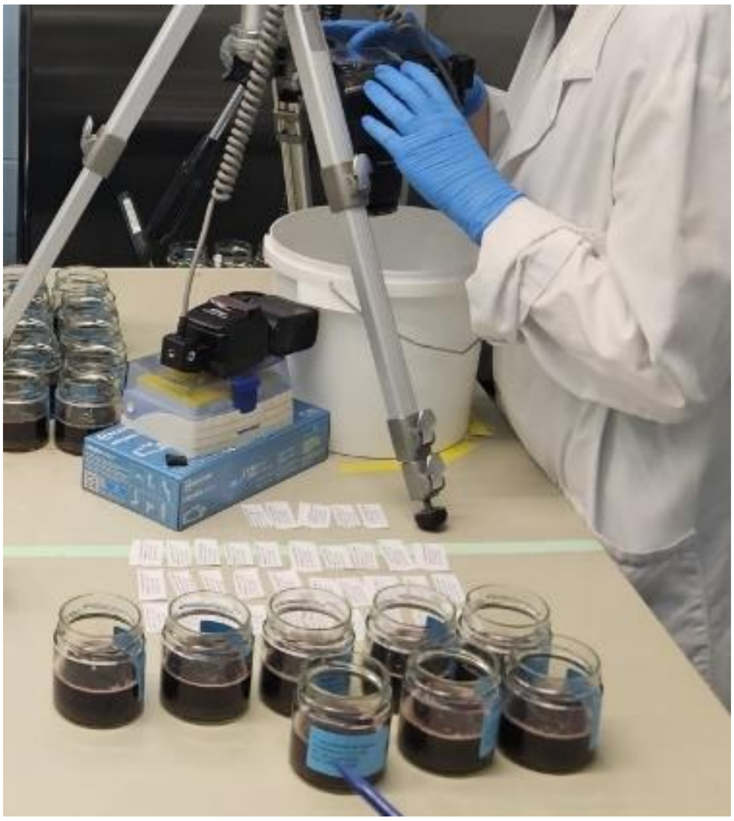

Supplementary Figure 1. Photography chamber for standardized imaging of OECD 232 test jars, constructed from a white bucket for light diffusion and equipped with a fixed-height tripod and ruler for scale.

#### Substrate

To enhance contrast, the water surface should be tinted with black ink, especially relevant for OECD artificial soils. Pelikan 4001 Brilliant Black proved effective without causing organism loss. A well-colored background improves model performance by distinguishing collembolans from foam, debris, or other interference (OECD, 2016).

#### Picture Acquisition & Counting

Once the water surface is still, test jar jars should be placed in the photography chamber for imaging (see figure 1). The photography chamber can consist of a white bucket to diffuse flash illumination and a tripod to maintain a stable and consistent camera distance. Each image should include a label with test details (name, treatment code, substance, date) and a ruler for scale. We recommend counting the adults at the time of picture acquisition (to prevent any adult counts from being missed) and to mark which test jars have <100 or 0 collembolans inside. Test jars with <100 animals should be validated manually after automated counting and those with 0 collembolans should not be fed to the YOLO pipeline. The previous step can be skipped if picture quality is very high but is still recommended. Manual counts for verification (recommended for low-quality images and/or test jars that do not contain visible Collembola, see below for more information) can be performed in e.g., ImageJ (Schneider et al., 2012) using the multi-point tool, placing markers at the center of each individual. Annotated images should be saved as TIFF files for ground-truth reference.

### Supplementary Information S3: AI Upscaling

#### GAN-based upscaling of low-resolution pictures (Coimbra)

To mitigate resolution limitations of the Coimbra dataset, we applied 2× super-resolution using Real-ESRGAN (X. Wang et al., 2021) as a preprocessing step prior to automated counting. Upscaling was performed on GPUs using FP16 inference and multi-GPU parallelization, with optional tiling to control memory usage. The procedure approximately quadruples the number of pixels per image, resulting in increased GPU memory requirements, runtime, and I/O overhead compared to direct inference on original images. Owing to this computational cost, upscaling was not considered a default preprocessing step but was evaluated specifically for its impact on counting performance in low-resolution imagery. An example of the visual effect of upscaling is provided in Supplementary Figure 2.

#### Effect of GAN-based upscaling on counting performance

For the populated concentrations of the low-resolution Coimbra dataset (C0–C2), GAN-based 2x upscaling led to a pronounced and statistically significant improvement in counting accuracy. Mean absolute counting errors (MAE) were reduced from MAE = 551.1 for original images to MAE = 80.7 for upscaled images, corresponding to an approximately 7-fold reduction in error. This improvement was highly significant under a paired Wilcoxon signed-rank test ($W = 120$, $p < 0.001$), demonstrating that upscaling consistently improved performance across replicates.

In contrast, at lower concentrations, upscaling increasingly amplified background texture and image noise, resulting in elevated false-positive counts. This indicates that the benefits of GAN-based super-resolution are density-dependent, and that indiscriminate application can be detrimental in sparse or near-empty OECD232 imagery. Accordingly, upscaling should be applied selectively, ideally restricted to images exceeding a minimum population density. While GAN-based upscaling entails substantial computational cost, it can be worthwhile for low-resolution datasets with dense populations, where it substantially improves object separability and counting accuracy. Examples of detection differences between original and upscaled images are provided in Supplementary Figure 2-3, together with a boxplot summarizing detection responses across concentration levels.

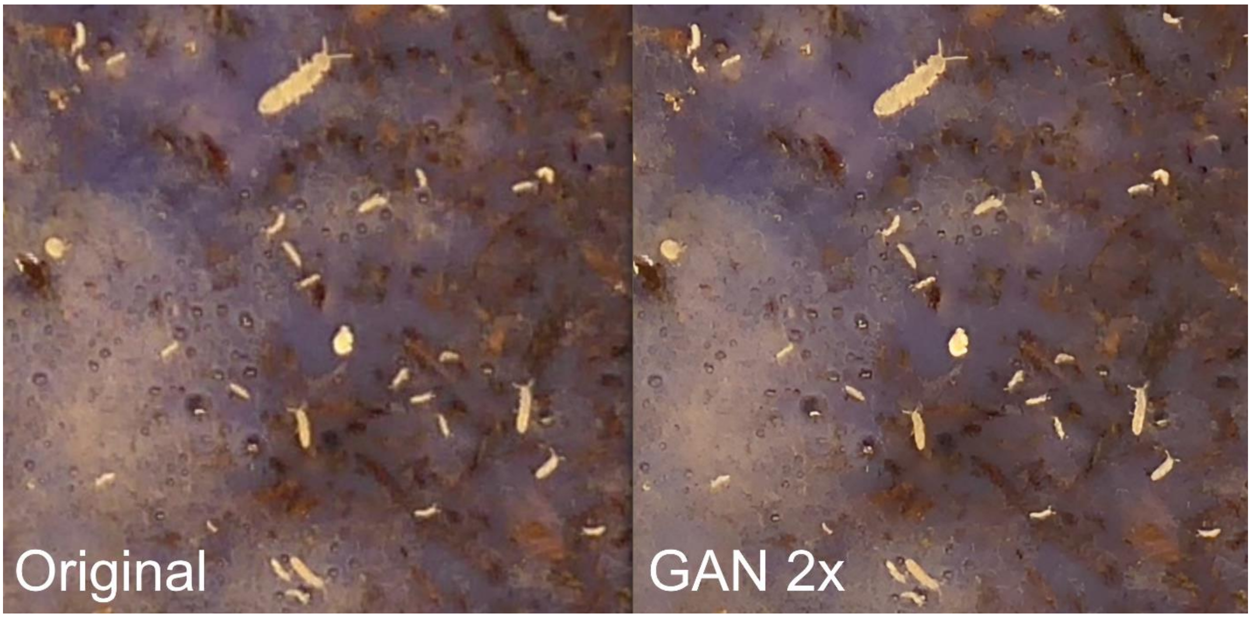

Supplementary Figure 2: The original panel on the left depicts a picture from the Coimbra dataset before the AI upscalind, while the GAN 2x picture on the right was after an AI upscaling (Wang et al., 2021). The AI upscaling was done to improve the quality of the picture to better detect springtails in low quality settings.

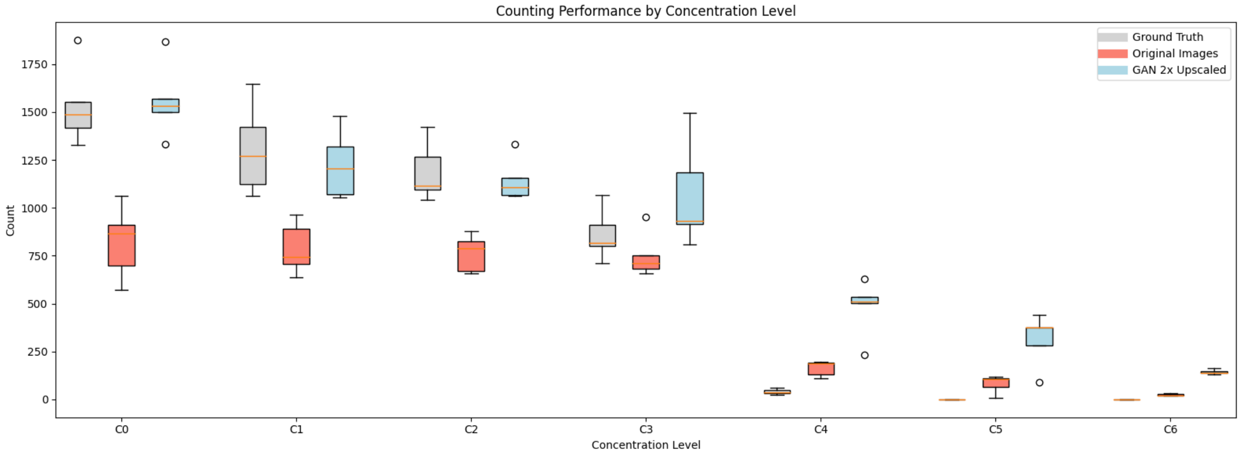

Supplementary Figure 3: Boxplot for the comparison of three different counting methods for the Coimbra dataset: Hand counted pictures serve as ground truth, Original Images counted with Collembot and GAN 2x upscaled images before counting with Collembot.

Supplementary Table 1. Overview of all experiments used for COLLEMBOT development and validation

| DATASET / EXPERIMENT | REFERENCE | SPECIES | SOIL TYPE(S) | CHEMICALS TESTED | CAMERA USED | NATIVE IMAGE RESOLUTION (PX) | CAMERA SETTINGS (ISO / APERTURE / FLASH) | PURPOSE | TRAINING TILES | EFFECT VALIDATION IMAGES |
|---|---|---|---|---|---|---|---|---|---|---|
| **BASEL** | Unpublished | *Folsomia candida* | LUFA 2.2 | Undisclosed | Nikon D800 + Speedlight flash | 6316 x 4912 | ISO 200 / f8 / Flash | Training + Validation | 417 tiles | 123 |
| **AMSTERDAM** | van Hall et al., 2025 | *Folsomia candida* | LUFA 2.2, OECD 2.5, 5, 10 | Chlorpyrifos, Cyproconazole, Imidacloprid, Lindane | Nikon Coolpix P510 | 4608 x 3456 | ISO 400 / f3.9 / No Flash | Independent Validation | 91 tiles | 1281 |
| **BAYREUTH** | Unpublished | *Folsomia candida* | OECD 10 | Polystyrene microplastic particles | Olympus E-M10 Mark II | 4608 x 3456 | ISO 5000 / f3.5 / No Flash | Training + Validation | 192 tiles | 56 |
| **COIMBRA** | Unpublished | *Folsomia candida* | LUFA 2.2 | Boric Acid | Panasonic Lumix CD-TZ90 | 3840x2160 | ISO 80-3200 / f3.3-6.4 / No Flash | Training + Validation | 27 tiles | 35 |
| **AARHUS (DENMARK)** | Wehrli et al., 2024 | *Folsomia candida* | LUFA 2.2 | Fluazinam | Sony ILCE-5000 | 4876 x 3632 | ISO 2000 / N/A / No Flash | Independent Validation | N/A | 209 |
| **COLLEMBOLAI** | Sys et al., 2022 | Mixed Collembola | Fluid Samples (no soil) | None | DSLR (APS-C & full-frame) on motorized rail + side flash Gloxy GX-F990C | Composites via stitching, e.g. 18’931×16’256 | ISO 100 / f11/ Flash | Training | 2,480 tiles | N/A |

### Supplementary Information S4:

Supplementary Table 2. YOLO v11 segXL performance metrics (F1-scores) for bounding boxes and masks across datasets as depicted in figure 4, with and without Flatbug CAI augmentation.

| DATASET | F1 (BBOX) WITH COLLEMBOLAI | F1 (MASK) WITH COLLEMBOLAI | F1 (BBOX) WITHOUT COLLEMBOLAI | F1 (MASK) WITHOUT COLLEMBOLAI |
|---|---|---|---|---|
| **AMSTERDAM** | 0.943 | 0.919 | 0.938 | 0.927 |
| **BAYREUTH** | 0.871 | 0.826 | 0.866 | 0.837 |
| **BASEL** | 0.938 | 0.933 | 0.946 | 0.939 |
| **COIMBRA** | 0.857 | 0.724 | 0.846 | 0.756 |
| **COLLEMBOLAI** | 0.88 | 0.805 | NA | NA |

Supplementary Table 3. Linear regression models comparing automated counts against manual counts of OECD 232 tests with *Folsomia candida,* for each dataset, including regression formula and coefficient of determination ($R^2$).

| SUBSTANCE | FORMULA | R2 |
|---|---|---|
| **AMSTERDAM_CHLORPYRIFOS_LUFA_2_2** | y = 0.9584x + 15.5929 | 0.955 |
| **AMSTERDAM_CHLORPYRIFOS_OECD10** | y = 1.0202x + 4.0228 | 0.979 |
| **AMSTERDAM_CHLORPYRIFOS_OECD2_5** | y = 1.0447x + 9.2442 | 0.984 |
| **AMSTERDAM_CHLORPYRIFOS_OECD5** | y = 0.943x + 20.8025 | 0.946 |
| **AMSTERDAM_CYPROCONAZOLE_OECD10** | y = 1.1198x + 30.6875 | 0.965 |
| **AMSTERDAM_CYPROCONAZOLE_OECD2_5** | y = 0.9993x + 7.8712 | 0.989 |
| **AMSTERDAM_CYPROCONAZOLE_OECD5** | y = 1.0146x + 4.821 | 0.969 |
| **AMSTERDAM_IMIDACLOPRID_LUFA_2_2** | y = 0.9638x + 11.3596 | 0.988 |
| **AMSTERDAM_IMIDACLOPRID_OECD10** | y = 1.0122x + 60.5752 | 0.904 |
| **AMSTERDAM_IMIDACLOPRID_OECD2_5** | y = 1.049x + 25.7469 | 0.978 |
| **AMSTERDAM_IMIDACLOPRID_OECD5** | y = 1.0459x + 13.7494 | 0.980 |
| **AMSTERDAM_LINDANE_LUFA_2_2** | y = 0.9184x - 38.6308 | 0.934 |
| **AMSTERDAM_LINDANE_OECD10** | y = 0.7483x + 81.6245 | 0.791 |
| **AMSTERDAM_LINDANE_OECD5** | y = 1.0113x - 3.8827 | 0.953 |
| **BASEL_LUFA_2_2** | y = 0.9631x - 2.4662 | 0.996 |
| **BAYREUTH_POLYSTYRENE_OECD_10** | y = 0.9068x + 60.9984 | 0.968 |

| | | |
|---|---|---|
| **COIMBRA_BORIC_ACID_OECD_5** | y = 0.9834x + 27.784 | 0.955 |
| **DENMARK_FLUAZINAM20_LUFA_2_2** | y = 0.8682x + 48.2077 | 0.902 |
| **DENMARK_FLUAZINAM22_LUFA_2_2** | y = 0.8398x + 67.8989 | 0.957 |
| **DENMARK_FLUAZINAM24_LUFA_2_2** | y = 0.846x + 48.6351 | 0.958 |
| **DENMARK_FLUAZINAM26_LUFA_2_2** | y = 0.9002x + 54.8671 | 0.958 |

Supplementary Table 4. Effect concentration estimates (EC10–EC90) for manual and automated counts of Collembola across substances, including confidence intervals.

| SUBSTANCE | METHOD | MODEL | EFFECT | EFFECT CONCENTRATION (MG $KG^{-1}$ DRY SOIL) | LOWER | UPPER |
|---|---|---|---|---|---|---|
| **BASEL_LUFA_2_2** | Automated | W1.3 | EC10 | 0.04 | -0.18 | 0.26 |
| **BASEL_LUFA_2_2** | Automated | W1.3 | EC20 | 0.19 | -0.57 | 0.94 |
| **BASEL_LUFA_2_2** | Automated | W1.3 | EC50 | 1.98 | -1.93 | 5.89 |
| **BASEL_LUFA_2_2** | Automated | W1.3 | EC90 | 24.1 | -23.2 | 71.4 |
| **BASEL_LUFA_2_2** | Manual | W1.3 | EC10 | 0.06 | -0.23 | 0.34 |
| **BASEL_LUFA_2_2** | Manual | W1.3 | EC20 | 0.25 | -0.69 | 1.18 |
| **BASEL_LUFA_2_2** | Manual | W1.3 | EC50 | 2.37 | -2.04 | 6.78 |
| **BASEL_LUFA_2_2** | Manual | W1.3 | EC90 | 25.9 | -22.48 | 74.3 |
| **BORIC_ACID_OECD5** | Automated | W1.3 | EC10 | 48.7 | 31.20 | 66.1 |
| **BORIC_ACID_OECD5** | Automated | W1.3 | EC20 | 60.1 | 44.5 | 75.7 |
| **BORIC_ACID_OECD5** | Automated | W1.3 | EC50 | 82.7 | 70.1 | 95.2 |
| **BORIC_ACID_OECD5** | Automated | W1.3 | EC90 | 115.9 | 92.6 | 139 |
| **BORIC_ACID_OECD5** | Manual | W1.3 | EC10 | 28.1 | 15 | 41.2 |
| **BORIC_ACID_OECD5** | Manual | W1.3 | EC20 | 39.8 | 26.7 | 53 |
| **BORIC_ACID_OECD5** | Manual | W1.3 | EC50 | 67.4 | 57.2 | 77.6 |
| **BORIC_ACID_OECD5** | Manual | W1.3 | EC90 | 117.8 | 96.1 | 139 |
| **CHLORPYRIFOS_LUFA_2_2** | Automated | W1.3 | EC10 | 0.10 | 0.07 | 0.12 |
| **CHLORPYRIFOS_LUFA_2_2** | Automated | W1.3 | EC20 | 0.11 | 0.09 | 0.13 |
| **CHLORPYRIFOS_LUFA_2_2** | Automated | W1.3 | EC50 | 0.13 | 0.11 | 0.15 |
| **CHLORPYRIFOS_LUFA_2_2** | Automated | W1.3 | EC90 | 0.16 | 0.14 | 0.17 |
| **CHLORPYRIFOS_LUFA_2_2** | Manual | W1.3 | EC10 | 0.11 | 0.04 | 0.18 |

| | | | | | | |
|---|---|---|---|---|---|---|
| **CHLORPYRIFOS_LUFA_2_2** | Manual | W1.3 | EC20 | 0.12 | 0.06 | 0.18 |
| **CHLORPYRIFOS_LUFA_2_2** | Manual | W1.3 | EC50 | 0.14 | 0.10 | 0.18 |
| **CHLORPYRIFOS_LUFA_2_2** | Manual | W1.3 | EC90 | 0.16 | 0.14 | 0.17 |
| **CHLORPYRIFOS_OECD10** | Automated | W1.3 | EC10 | 0.13 | 0.00 | 0.26 |
| **CHLORPYRIFOS_OECD10** | Automated | W1.3 | EC20 | 0.14 | 0.03 | 0.24 |
| **CHLORPYRIFOS_OECD10** | Automated | W1.3 | EC50 | 0.15 | 0.08 | 0.21 |
| **CHLORPYRIFOS_OECD10** | Automated | W1.3 | EC90 | 0.16 | 0.13 | 0.18 |
| **CHLORPYRIFOS_OECD10** | Manual | W1.3 | EC10 | 0.10 | 0.08 | 0.12 |
| **CHLORPYRIFOS_OECD10** | Manual | W1.3 | EC20 | 0.11 | 0.09 | 0.13 |
| **CHLORPYRIFOS_OECD10** | Manual | W1.3 | EC50 | 0.13 | 0.11 | 0.14 |
| **CHLORPYRIFOS_OECD10** | Manual | W1.3 | EC90 | 0.15 | 0.13 | 0.17 |
| **CHLORPYRIFOS_OECD2_5** | Automated | W1.3 | EC10 | 0.03 | 0.00 | 0.06 |
| **CHLORPYRIFOS_OECD2_5** | Automated | W1.3 | EC20 | 0.03 | 0.01 | 0.05 |
| **CHLORPYRIFOS_OECD2_5** | Automated | W1.3 | EC50 | 0.04 | 0.03 | 0.05 |
| **CHLORPYRIFOS_OECD2_5** | Automated | W1.3 | EC90 | 0.04 | 0.03 | 0.06 |
| **CHLORPYRIFOS_OECD2_5** | Manual | W1.3 | EC10 | 0.03 | 0.03 | 0.04 |
| **CHLORPYRIFOS_OECD2_5** | Manual | W1.3 | EC20 | 0.04 | 0.03 | 0.04 |
| **CHLORPYRIFOS_OECD2_5** | Manual | W1.3 | EC50 | 0.04 | 0.04 | 0.04 |
| **CHLORPYRIFOS_OECD2_5** | Manual | W1.3 | EC90 | 0.04 | 0.04 | 0.04 |
| **CHLORPYRIFOS_OECD5** | Automated | W1.3 | EC10 | 0.07 | 0.01 | 0.12 |
| **CHLORPYRIFOS_OECD5** | Automated | W1.3 | EC20 | 0.07 | 0.03 | 0.11 |
| **CHLORPYRIFOS_OECD5** | Automated | W1.3 | EC50 | 0.08 | 0.06 | 0.10 |
| **CHLORPYRIFOS_OECD5** | Automated | W1.3 | EC90 | 0.08 | 0.08 | 0.09 |
| **CHLORPYRIFOS_OECD5** | Manual | W1.3 | EC10 | 0.06 | 0.01 | 0.12 |
| **CHLORPYRIFOS_OECD5** | Manual | W1.3 | EC20 | 0.07 | 0.02 | 0.11 |
| **CHLORPYRIFOS_OECD5** | Manual | W1.3 | EC50 | 0.07 | 0.05 | 0.10 |
| **CHLORPYRIFOS_OECD5** | Manual | W1.3 | EC90 | 0.08 | 0.07 | 0.09 |
| **CYPROCONAZOLE_OECD10** | Automated | W1.3 | EC10 | 155 | 68.7 | 242 |
| **CYPROCONAZOLE_OECD10** | Automated | W1.3 | EC20 | 244 | 149 | 341 |
| **CYPROCONAZOLE_OECD10** | Automated | W1.3 | EC50 | 485 | 392 | 579 |
| **CYPROCONAZOLE_OECD10** | Automated | W1.3 | EC90 | 1004 | 729 | 1279 |

| | | | | | | |
|---|---|---|---|---|---|---|
| **CYPROCONAZOLE_OECD10** | Manual | W1.3 | EC10 | 184 | 75.2 | 293 |
| **CYPROCONAZOLE_OECD10** | Manual | W1.3 | EC20 | 275 | 161 | 389 |
| **CYPROCONAZOLE_OECD10** | Manual | W1.3 | EC50 | 503 | 403 | 603 |
| **CYPROCONAZOLE_OECD10** | Manual | W1.3 | EC90 | 953 | 683 | 1223 |
| **CYPROCONAZOLE_OECD2_5** | Automated | W1.3 | EC10 | 36.1 | 17.6 | 54.6 |
| **CYPROCONAZOLE_OECD2_5** | Automated | W1.3 | EC20 | 55.2 | 35.4 | 75.1 |
| **CYPROCONAZOLE_OECD2_5** | Automated | W1.3 | EC50 | 105 | 83.1 | 127 |
| **CYPROCONAZOLE_OECD2_5** | Automated | W1.3 | EC90 | 207 | 140 | 274 |
| **CYPROCONAZOLE_OECD2_5** | Manual | W1.3 | EC10 | 37.2 | 15.4 | 58.9 |
| **CYPROCONAZOLE_OECD2_5** | Manual | W1.3 | EC20 | 54.6 | 33.7 | 75.5 |
| **CYPROCONAZOLE_OECD2_5** | Manual | W1.3 | EC50 | 97.5 | 77.8 | 117 |
| **CYPROCONAZOLE_OECD2_5** | Manual | W1.3 | EC90 | 180.3 | 103.74 | 256.81 |
| **CYPROCONAZOLE_OECD5** | Automated | W1.3 | EC10 | 47.7 | 16.0 | 79.5 |
| **CYPROCONAZOLE_OECD5** | Automated | W1.3 | EC20 | 81.9 | 43.2 | 121 |
| **CYPROCONAZOLE_OECD5** | Automated | W1.3 | EC50 | 185 | 135 | 236 |
| **CYPROCONAZOLE_OECD5** | Automated | W1.3 | EC90 | 440 | 268 | 611 |
| **CYPROCONAZOLE_OECD5** | Manual | W1.3 | EC10 | 51.2 | 20.8 | 81.7 |
| **CYPROCONAZOLE_OECD5** | Manual | W1.3 | EC20 | 85.6 | 49.4 | 122 |
| **CYPROCONAZOLE_OECD5** | Manual | W1.3 | EC50 | 186 | 140 | 232 |
| **CYPROCONAZOLE_OECD5** | Manual | W1.3 | EC90 | 422 | 272 | 572 |
| **FLUAZINAM20_LUFA_2_2** | Automated | W1.3 | EC10 | 7.90 | 5.22 | 10.6 |
| **FLUAZINAM20_LUFA_2_2** | Automated | W1.3 | EC20 | 9.22 | 6.94 | 11.51 |
| **FLUAZINAM20_LUFA_2_2** | Automated | W1.3 | EC50 | 11.7 | 9.47 | 13.8 |
| **FLUAZINAM20_LUFA_2_2** | Automated | W1.3 | EC90 | 14.9 | 10.7 | 19.1 |
| **FLUAZINAM20_LUFA_2_2** | Manual | W1.3 | EC10 | 6.11 | 2.42 | 9.80 |
| **FLUAZINAM20_LUFA_2_2** | Manual | W1.3 | EC20 | 7.57 | 4.47 | 10.7 |
| **FLUAZINAM20_LUFA_2_2** | Manual | W1.3 | EC50 | 10.5 | 8.22 | 12.7 |
| **FLUAZINAM20_LUFA_2_2** | Manual | W1.3 | EC90 | 14.8 | 9.05 | 20.5 |
| **FLUAZINAM22_LUFA_2_2** | Automated | W1.3 | EC10 | 5.41 | 1.41 | 9.41 |
| **FLUAZINAM22_LUFA_2_2** | Automated | W1.3 | EC20 | 7.11 | 3.31 | 10.9 |
| **FLUAZINAM22_LUFA_2_2** | Automated | W1.3 | EC50 | 10.7 | 7.37 | 14.1 |

| | | | | | | |
|---|---|---|---|---|---|---|
| **FLUAZINAM22_LUFA_2_2** | Automated | W1.3 | EC90 | 16.7 | 9.84 | 23.5 |
| **FLUAZINAM22_LUFA_2_2** | Manual | W1.3 | EC10 | 4.26 | 0.38 | 8.15 |
| **FLUAZINAM22_LUFA_2_2** | Manual | W1.3 | EC20 | 5.85 | 2.03 | 9.68 |
| **FLUAZINAM22_LUFA_2_2** | Manual | W1.3 | EC50 | 9.46 | 5.54 | 13.37 |
| **FLUAZINAM22_LUFA_2_2** | Manual | W1.3 | EC90 | 15.7 | 6.21 | 25.22 |
| **FLUAZINAM24_LUFA_2_2** | Automated | W1.3 | EC10 | 6.48 | 2.53 | 10.43 |
| **FLUAZINAM24_LUFA_2_2** | Automated | W1.3 | EC20 | 7.87 | 4.60 | 11.13 |
| **FLUAZINAM24_LUFA_2_2** | Automated | W1.3 | EC50 | 10.5 | 8.06 | 13.04 |
| **FLUAZINAM24_LUFA_2_2** | Automated | W1.3 | EC90 | 14.4 | 8.38 | 20.43 |
| **FLUAZINAM24_LUFA_2_2** | Manual | W1.3 | EC10 | 4.98 | 1.19 | 8.76 |
| **FLUAZINAM24_LUFA_2_2** | Manual | W1.3 | EC20 | 6.39 | 2.99 | 9.80 |
| **FLUAZINAM24_LUFA_2_2** | Manual | W1.3 | EC50 | 9.34 | 6.66 | 12.03 |
| **FLUAZINAM24_LUFA_2_2** | Manual | W1.3 | EC90 | 14.0 | 7.79 | 20.13 |
| **FLUAZINAM26_LUFA_2_2** | Automated | W1.3 | EC10 | 3.80 | 0.07 | 7.52 |
| **FLUAZINAM26_LUFA_2_2** | Automated | W1.3 | EC20 | 5.32 | 1.65 | 8.99 |
| **FLUAZINAM26_LUFA_2_2** | Automated | W1.3 | EC50 | 8.85 | 6.10 | 11.61 |
| **FLUAZINAM26_LUFA_2_2** | Automated | W1.3 | EC90 | 15.2 | 9.14 | 21.21 |
| **FLUAZINAM26_LUFA_2_2** | Manual | W1.3 | EC10 | 2.68 | -0.77 | 6.13 |
| **FLUAZINAM26_LUFA_2_2** | Manual | W1.3 | EC20 | 3.98 | 0.41 | 7.54 |
| **FLUAZINAM26_LUFA_2_2** | Manual | W1.3 | EC50 | 7.23 | 4.43 | 10.03 |
| **FLUAZINAM26_LUFA_2_2** | Manual | W1.3 | EC90 | 13.6 | 6.17 | 21.07 |
| **IMIDACLOPRID_LUFA_2_2** | Automated | W1.3 | EC10 | 0.29 | 0.21 | 0.37 |
| **IMIDACLOPRID_LUFA_2_2** | Automated | W1.3 | EC20 | 0.39 | 0.31 | 0.48 |
| **IMIDACLOPRID_LUFA_2_2** | Automated | W1.3 | EC50 | 0.64 | 0.54 | 0.73 |
| **IMIDACLOPRID_LUFA_2_2** | Automated | W1.3 | EC90 | 1.06 | 0.85 | 1.26 |
| **IMIDACLOPRID_LUFA_2_2** | Manual | W1.3 | EC10 | 0.24 | 0.18 | 0.31 |
| **IMIDACLOPRID_LUFA_2_2** | Manual | W1.3 | EC20 | 0.34 | 0.28 | 0.41 |
| **IMIDACLOPRID_LUFA_2_2** | Manual | W1.3 | EC50 | 0.57 | 0.49 | 0.65 |
| **IMIDACLOPRID_LUFA_2_2** | Manual | W1.3 | EC90 | 0.99 | 0.77 | 1.20 |
| **IMIDACLOPRID_OECD10** | Automated | W1.3 | EC10 | 2.49 | -0.37 | 5.35 |
| **IMIDACLOPRID_OECD10** | Automated | W1.3 | EC20 | 2.77 | 0.57 | 4.96 |

| | | | | | | |
|---|---|---|---|---|---|---|
| **IMIDACLOPRID_OECD10** | Automated | W1.3 | EC50 | 3.25 | 2.41 | 4.09 |
| **IMIDACLOPRID_OECD10** | Automated | W1.3 | EC90 | 3.86 | 2.60 | 5.12 |
| **IMIDACLOPRID_OECD10** | Manual | W1.3 | EC10 | 0.68 | 0.18 | 1.18 |
| **IMIDACLOPRID_OECD10** | Manual | W1.3 | EC20 | 1.07 | 0.51 | 1.63 |
| **IMIDACLOPRID_OECD10** | Manual | W1.3 | EC50 | 2.12 | 1.58 | 2.66 |
| **IMIDACLOPRID_OECD10** | Manual | W1.3 | EC90 | 4.39 | 3.00 | 5.79 |
| **IMIDACLOPRID_OECD2_5** | Automated | W1.3 | EC10 | 0.60 | 0.26 | 0.94 |
| **IMIDACLOPRID_OECD2_5** | Automated | W1.3 | EC20 | 0.74 | 0.45 | 1.03 |
| **IMIDACLOPRID_OECD2_5** | Automated | W1.3 | EC50 | 1.02 | 0.88 | 1.15 |
| **IMIDACLOPRID_OECD2_5** | Automated | W1.3 | EC90 | 1.42 | 1.18 | 1.66 |
| **IMIDACLOPRID_OECD2_5** | Manual | W1.3 | EC10 | 0.35 | 0.24 | 0.45 |
| **IMIDACLOPRID_OECD2_5** | Manual | W1.3 | EC20 | 0.49 | 0.38 | 0.60 |
| **IMIDACLOPRID_OECD2_5** | Manual | W1.3 | EC50 | 0.82 | 0.73 | 0.92 |
| **IMIDACLOPRID_OECD2_5** | Manual | W1.3 | EC90 | 1.43 | 1.26 | 1.60 |
| **IMIDACLOPRID_OECD5** | Automated | W1.3 | EC10 | 0.80 | 0.45 | 1.16 |
| **IMIDACLOPRID_OECD5** | Automated | W1.3 | EC20 | 1.11 | 0.74 | 1.47 |
| **IMIDACLOPRID_OECD5** | Automated | W1.3 | EC50 | 1.81 | 1.30 | 2.31 |
| **IMIDACLOPRID_OECD5** | Automated | W1.3 | EC90 | 3.03 | 1.76 | 4.31 |
| **IMIDACLOPRID_OECD5** | Manual | W1.3 | EC10 | 0.57 | 0.25 | 0.89 |
| **IMIDACLOPRID_OECD5** | Manual | W1.3 | EC20 | 0.85 | 0.52 | 1.18 |
| **IMIDACLOPRID_OECD5** | Manual | W1.3 | EC50 | 1.56 | 1.15 | 1.97 |
| **IMIDACLOPRID_OECD5** | Manual | W1.3 | EC90 | 2.97 | 1.62 | 4.33 |
| **LINDANE_LUFA_2_2** | Automated | W1.3 | EC10 | 0.44 | -0.03 | 0.90 |
| **LINDANE_LUFA_2_2** | Automated | W1.3 | EC20 | 0.66 | 0.18 | 1.14 |
| **LINDANE_LUFA_2_2** | Automated | W1.3 | EC50 | 1.24 | 0.84 | 1.64 |
| **LINDANE_LUFA_2_2** | Automated | W1.3 | EC90 | 2.41 | 1.12 | 3.69 |
| **LINDANE_LUFA_2_2** | Manual | W1.3 | EC10 | 0.70 | 0.25 | 1.15 |
| **LINDANE_LUFA_2_2** | Manual | W1.3 | EC20 | 0.95 | 0.53 | 1.38 |
| **LINDANE_LUFA_2_2** | Manual | W1.3 | EC50 | 1.52 | 1.19 | 1.85 |
| **LINDANE_LUFA_2_2** | Manual | W1.3 | EC90 | 2.49 | 1.63 | 3.36 |
| **LINDANE_OECD10** | Automated | W1.3 | EC10 | 1.26 | 0.31 | 2.21 |

| | | | | | | |
|---|---|---|---|---|---|---|
| **LINDANE_OECD10** | Automated | W1.3 | EC20 | 1.72 | 0.82 | 2.61 |
| **LINDANE_OECD10** | Automated | W1.3 | EC50 | 2.74 | 2.13 | 3.35 |
| **LINDANE_OECD10** | Automated | W1.3 | EC90 | 4.49 | 3.08 | 5.91 |
| **LINDANE_OECD10** | Manual | W1.3 | EC10 | 0.77 | 0.47 | 1.07 |
| **LINDANE_OECD10** | Manual | W1.3 | EC20 | 1.15 | 0.84 | 1.47 |
| **LINDANE_OECD10** | Manual | W1.3 | EC50 | 2.11 | 1.80 | 2.43 |
| **LINDANE_OECD10** | Manual | W1.3 | EC90 | 4.01 | 3.16 | 4.87 |
| **LINDANE_OECD5** | Automated | W1.3 | EC10 | 0.13 | 0.00 | 0.26 |
| **LINDANE_OECD5** | Automated | W1.3 | EC20 | 0.29 | 0.08 | 0.49 |
| **LINDANE_OECD5** | Automated | W1.3 | EC50 | 0.96 | 0.63 | 1.29 |
| **LINDANE_OECD5** | Automated | W1.3 | EC90 | 3.49 | 2.06 | 4.91 |
| **LINDANE_OECD5** | Manual | W1.3 | EC10 | 0.23 | 0.06 | 0.40 |
| **LINDANE_OECD5** | Manual | W1.3 | EC20 | 0.43 | 0.20 | 0.66 |
| **LINDANE_OECD5** | Manual | W1.3 | EC50 | 1.13 | 0.84 | 1.42 |
| **LINDANE_OECD5** | Manual | W1.3 | EC90 | 3.13 | 2.11 | 4.16 |

#### Supplementary Information S5: NOEC/LOEC detection

For each substance and method (Manual and Automated), data were analyzed as follows:

- **Assumption Checks:** Normality of residuals was tested using the Shapiro–Wilk test and homogeneity of variances using Levene's test.
- **Parametric Approach:** If assumptions were met, one-way ANOVA was applied followed by Dunnett's post-hoc test to compare each concentration against the control.
- **Non-Parametric Approach:** If assumptions failed, Kruskal–Wallis test was used, followed by Dunn's post-hoc test with Bonferroni adjustment.
- **NOEC and LOEC Determination:**
  - NOEC (No Observed Effect Concentration) was defined as the highest tested concentration without a statistically significant difference from the control.

- LOEC (Lowest Observed Effect Concentration) was defined as the lowest tested concentration with a statistically significant difference from the control.
  Both values were calculated by sorting concentrations in ascending order and applying significance thresholds (α = 0.05).

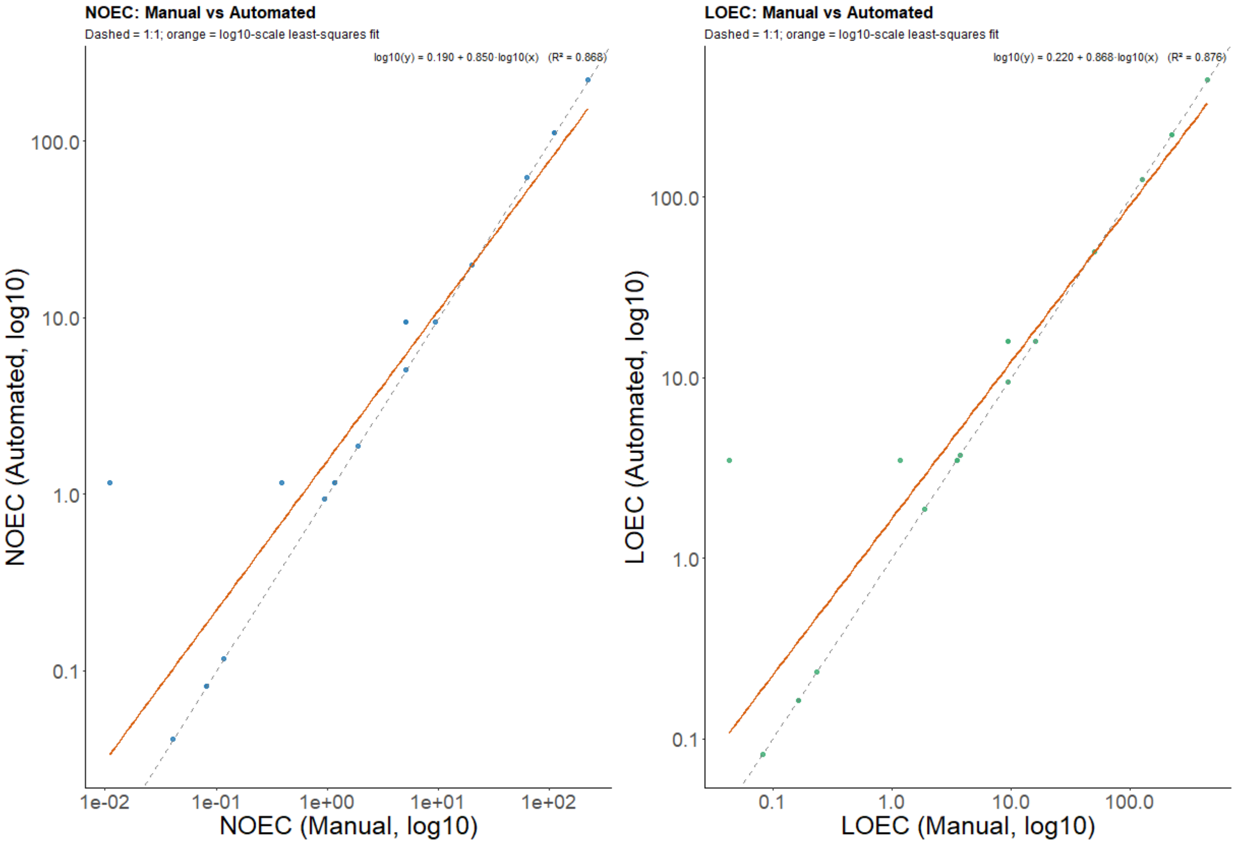

Supplementary Figure 4: Comparison of NOEC and LOEC values for the toxicity of pesticides to *Folsomia candida*, obtained from automated versus manual counts. The dotted line represents the 1:1 identity line, while the solid orange line shows the log10-scale east squares regression fit between the two methods. Regression equations and $R^2$ values are displayed for each panel.

Supplementary Table 5. Summary of NOEC and LOEC values (mg kg$^{-1}$ dry soil) for all tested substances and counting methods (Manual vs Automated), including the statistical approach applied (ANOVA + Dunnett or Kruskal–Wallis + Dunn). NOEC represents the highest concentration without a statistically significant effect compared to the control and LOEC represents the lowest concentration with a significant effect (α = 0.05). All concentrations are rounded to three decimal places.

| SUBSTANCE | ENDPOINT | METHOD | NOEC | LOEC | P AT LOEC | SHAPIRO P | LEVENE P | N |
|---|---|---|---|---|---|---|---|---|
| **AMSTERDAM_CHLORPYRIFOS_LUFA_2_2** | Automated | ANOVA + Dunnett (vs. control) | 0.082 | 0.164 | 0.000 | 0.560 | 0.062 | 40 |
| **AMSTERDAM_CHLORPYRIFOS_LUFA_2_2** | Manual | ANOVA + Dunnett (vs. control) | 0.082 | 0.164 | 0.000 | 0.548 | 0.125 | 40 |
| **AMSTERDAM_CHLORPYRIFOS_OECD10** | Automated | Kruskal–Wallis + Dunn (Holm, vs. control) | 0.082 | 0.164 | 0.014 | 0.009 | 0.223 | 40 |
| **AMSTERDAM_CHLORPYRIFOS_OECD10** | Manual | Kruskal–Wallis + Dunn (Holm, vs. control) | 0.082 | 0.164 | 0.007 | 0.009 | 0.028 | 40 |
| **AMSTERDAM_CHLORPYRIFOS_OECD2_5** | Automated | Kruskal–Wallis + Dunn (Holm, vs. control) | 0.041 | 0.082 | 0.000 | 0.000 | 0.001 | 43 |
| **AMSTERDAM_CHLORPYRIFOS_OECD2_5** | Manual | Kruskal–Wallis + Dunn (Holm, vs. control) | 0.041 | 0.082 | 0.004 | 0.013 | 0.023 | 39 |
| **AMSTERDAM_CHLORPYRIFOS_OECD5** | Automated | Kruskal–Wallis + Dunn (Holm, vs. control) | 0.082 | 0.164 | 0.002 | 0.227 | 0.019 | 41 |
| **AMSTERDAM_CHLORPYRIFOS_OECD5** | Manual | Kruskal–Wallis + Dunn (Holm, vs. control) | 0.082 | 0.164 | 0.004 | 0.063 | 0.010 | 40 |
| **AMSTERDAM_CYPROCONAZOLE_OECD10** | Automated | ANOVA + Dunnett (vs. control) | 222 | 445 | 0.000 | 0.777 | 0.250 | 40 |
| **AMSTERDAM_CYPROCONAZOLE_OECD10** | Manual | ANOVA + Dunnett (vs. control) | 222 | 445 | 0.000 | 0.116 | 0.163 | 40 |
| **AMSTERDAM_CYPROCONAZOLE_OECD2_5** | Automated | Kruskal–Wallis + Dunn (Holm, vs. control) | 111 | 222 | 0.011 | 0.001 | 0.086 | 40 |
| **AMSTERDAM_CYPROCONAZOLE_OECD2_5** | Manual | Kruskal–Wallis + Dunn (Holm, vs. control) | 111 | 222 | 0.009 | 0.002 | 0.184 | 40 |
| **AMSTERDAM_CYPROCONAZOLE_OECD5** | Automated | Kruskal–Wallis + Dunn (Holm, vs. control) | 111 | 222 | 0.047 | 0.000 | 0.048 | 40 |
| **AMSTERDAM_CYPROCONAZOLE_OECD5** | Manual | Kruskal–Wallis + Dunn (Holm, vs. control) | 111 | 222 | 0.044 | 0.038 | 0.018 | 40 |
| **AMSTERDAM_IMIDACLOPRID_LUFA_2_2** | Automated | Kruskal–Wallis + Dunn (Holm, vs. control) | 1.17 | 3.51 | 0.039 | 0.004 | 0.168 | 40 |
| **AMSTERDAM_IMIDACLOPRID_LUFA_2_2** | Manual | Kruskal–Wallis + Dunn (Holm, vs. control) | 1.17 | 3.51 | 0.004 | 0.005 | 0.041 | 40 |
| **AMSTERDAM_IMIDACLOPRID_OECD10** | Automated | ANOVA + Dunnett (vs. control) | 1.17 | 3.51 | 0.000 | 0.414 | 0.272 | 40 |
| **AMSTERDAM_IMIDACLOPRID_OECD10** | Manual | ANOVA + Dunnett (vs. control) | 0.011 | 0.043 | 0.049 | 0.450 | 0.210 | 40 |
| **AMSTERDAM_IMIDACLOPRID_OECD2_5** | Automated | Kruskal–Wallis + Dunn (Holm, vs. control) | 1.17 | 3.507 | 0.016 | 0.260 | 0.050 | 39 |
| **AMSTERDAM_IMIDACLOPRID_OECD2_5** | Manual | Kruskal–Wallis + Dunn (Holm, vs. control) | 0.39 | 1.17 | 0.025 | 0.001 | 0.029 | 39 |
| **AMSTERDAM_IMIDACLOPRID_OECD5** | Automated | Kruskal–Wallis + Dunn (Holm, vs. control) | 1.17 | 3.51 | 0.025 | 0.034 | 0.095 | 40 |
| **AMSTERDAM_IMIDACLOPRID_OECD5** | Manual | Kruskal–Wallis + Dunn (Holm, vs. control) | 1.17 | 3.51 | 0.026 | 0.014 | 0.207 | 40 |
| **AMSTERDAM_LINDANE_LUFA_2_2** | Automated | Kruskal–Wallis + Dunn (Holm, vs. control) | 0.936 | 1.87 | 0.008 | 0.007 | 0.005 | 40 |
| **AMSTERDAM_LINDANE_LUFA_2_2** | Manual | Kruskal–Wallis + Dunn (Holm, vs. control) | 0.936 | 1.87 | 0.038 | 0.017 | 0.079 | 40 |
| **AMSTERDAM_LINDANE_OECD10** | Automated | ANOVA + Dunnett (vs. control) | 1.87 | 3.74 | 0.000 | 0.332 | 0.632 | 40 |
| **AMSTERDAM_LINDANE_OECD10** | Manual | Kruskal–Wallis + Dunn (Holm, vs. control) | 1.87 | 3.74 | 0.006 | 0.942 | 0.040 | 40 |

| | | | | | | | | |
|---|---|---|---|---|---|---|---|---|
| **AMSTERDAM_LINDANE_OECD5** | Automated | ANOVA + Dunnett (vs. control) | 0.117 | 0.23 | 0.001 | 0.973 | 0.155 | 39 |
| **AMSTERDAM_LINDANE_OECD5** | Manual | ANOVA + Dunnett (vs. control) | 0.117 | 0.23 | 0.031 | 0.405 | 0.464 | 39 |
| **BASEL_LUFA_2_2** | Automated | Kruskal–Wallis + Dunn (Holm, vs. control) | 20 | 50 | 0.007 | 0.017 | 0.009 | 26 |
| **BASEL_LUFA_2_2** | Manual | Kruskal–Wallis + Dunn (Holm, vs. control) | 20 | 50 | 0.007 | 0.022 | 0.028 | 26 |
| **COIMBRA_BORIC_ACID_OECD_5** | Automated | Kruskal–Wallis + Dunn (Holm, vs. control) | 62.5 | 125 | 0.009 | 0.001 | 0.073 | 35 |
| **COIMBRA_BORIC_ACID_OECD_5** | Manual | Kruskal–Wallis + Dunn (Holm, vs. control) | 62.5 | 125 | 0.016 | 0.002 | 0.034 | 35 |
| **DENMARK_FLUAZINAM20_LUFA_2_2** | Automated | Kruskal–Wallis + Dunn (Holm, vs. control) | 9.440 | 15.981 | 0.006 | 0.843 | 0.004 | 42 |
| **DENMARK_FLUAZINAM20_LUFA_2_2** | Manual | ANOVA + Dunnett (vs. control) | 5.060 | 9.440 | 0.009 | 0.665 | 0.377 | 42 |
| **DENMARK_FLUAZINAM22_LUFA_2_2** | Automated | ANOVA + Dunnett (vs. control) | 9.440 | 15.981 | 0.000 | 0.520 | 0.215 | 43 |
| **DENMARK_FLUAZINAM22_LUFA_2_2** | Manual | ANOVA + Dunnett (vs. control) | 5.060 | 9.440 | 0.022 | 0.532 | 0.353 | 43 |
| **DENMARK_FLUAZINAM24_LUFA_2_2** | Automated | ANOVA + Dunnett (vs. control) | 9.440 | 15.981 | 0.000 | 0.264 | 0.179 | 44 |
| **DENMARK_FLUAZINAM24_LUFA_2_2** | Manual | ANOVA + Dunnett (vs. control) | 9.440 | 15.981 | 0.000 | 0.236 | 0.126 | 44 |
| **DENMARK_FLUAZINAM26_LUFA_2_2** | Automated | Kruskal–Wallis + Dunn (Holm, vs. control) | 5.060 | 9.440 | 0.036 | 0.003 | 0.555 | 44 |
| **DENMARK_FLUAZINAM26_LUFA_2_2** | Manual | Kruskal–Wallis + Dunn (Holm, vs. control) | 5.060 | 9.440 | 0.016 | 0.003 | 0.472 | 44 |
